# Ecological processes shaping marine microbial assemblages diverge between equatorial and temperate time-series

**DOI:** 10.1101/2025.08.21.671554

**Authors:** Pedro C. Junger, Vinicius S. Kavagutti, Ina Maria Deutschmann, Carlota R. Gazulla, Paula Huber, Maiara Menezes, Rodolfo Paranhos, André M. Amado, Isabel Ferrera, Janaina Rigonato, Samuel Chaffron, Josep M. Gasol, Ramiro Logares, Hugo Sarmento

## Abstract

Marine microbial communities are structured by a complex interplay of deterministic and stochastic processes, yet how these vary across latitudes remains poorly understood. Most long-term microbial observatories are restricted to temperate regions, limiting our ability to assess latitudinal contrasts in microbial dynamics. Here, we compare coastal microbial communities from two contrasting marine time-series stations using standardized molecular protocols: a new tropical site in the Equatorial Atlantic (EAMO, 6°S) and a well-studied temperate site in the Mediterranean Sea (BBMO, 41°N). Monthly 16S and 18S rRNA gene sequencing of two size-fractions (0.22–3 µm and >3 µm) over 41 months (from April 2013 to August 2016) revealed marked differences in taxonomic composition, temporal variability, and ecological assembly processes. Temperate communities exhibited strong seasonal turnover, higher beta-diversity, and tighter coupling with environmental variables such as temperature and daylength. In contrast, tropical communities were compositionally more stable and more governed by biotic factors and stochastic processes such as historical contingency and ecological drift. These patterns were consistent across taxonomic domains and size-fractions, though selection was generally stronger in prokaryotes and the smallest size-fraction. Co-occurrence networks at the temperate site were more densely connected and environmentally responsive compared to tropical networks, where stochastic processes and putative biological interactions gain prominence. This study highlights the importance of integrating observatories from underrepresented latitudes into global microbial monitoring efforts, particularly as climate change alters the amplitude and frequency of environmental drivers across the ocean.

## Introduction

Marine microbial communities constitute a substantial component of Earth’s biodiversity (de Vargas et al., 2015; Locey & Lennon, 2016; McNichol et al., 2025), and their interactions are fundamental for Earth’s ecosystem functioning and biogeochemical cycles (Falkowski et al., 2008; Guidi et al., 2016). Understanding the factors and mechanisms that drive microbial dynamics is important to predict how ongoing environmental changes impact their community composition, interactions, and functions (Chaffron et al., 2021; Doney et al., 2012). Global oceanographic expeditions (e.g., *Tara* Oceans and Malaspina), and ocean-basin transects (e.g., BioGEOTRACES and Bio-GO-SHIP) have revealed large-scale spatial patterns of microbial diversity (Giner et al., 2020; Ibarbalz et al., 2019; McNichol et al., 2025; Salazar et al., 2016; Sunagawa et al., 2015), and their ecological associations (Chaffron et al., 2021; Deutschmann et al., 2024). The ecological processes shaping microbial communities have been shown to change between biological domains (prokaryotes vs. eukaryotes), as well as with ocean depth and spatial scale (Junger et al., 2023; Logares et al., 2020). However, these space-for-time studies lack the temporal dimension, which is essential to fully understand the patterns of microbial diversity and the underlying mechanisms shaping ocean microbial communities (Buttigieg et al., 2018; Moreira & López-García, 2019).

Most long-term microbial observatories that have collected molecular data remain concentrated in northern mid-latitudes (∼22° to 50°) and often lack methodological standardization (e.g., primers pairs, sequencing technology), limiting their inter-comparison (Buttigieg et al., 2018; Raes et al., 2024). Meanwhile, most of the world’s ocean surface is under warm oligotrophic tropical conditions (Behrenfeld et al., 2006), where primary production is usually dominated by picoplankton (<2 µm), including prokaryotes (*Synechococcus* and *Prochlorococcus*) and small single-cell eukaryotes (Li et al., 1983; Platt et al., 1983; Tarran et al., 2006). In this context, comprehensive comparative studies of ocean time-series using standardized methods to assess microbial diversity and associations in the low-latitude regions remain scarce, especially considering understudied areas such as the South Atlantic Ocean. There are some remarkable exceptions, though, such as the Australian Microbiome Initiative, that generated a dataset – conceived with standard methods and protocols – of seven coastal microbial time-series covering a latitudinal gradient (∼12° to 42°S) at continental scale (Brown et al., 2018). More recently, among these time-series, the tropical Yongala observatory (∼19°S) was shown to exhibit weak seasonal recurrence and increasing community dissimilarity over time, in contrast to several higher-latitude sites, suggesting more stability and drift in tropical marine microbial communities (Raes et al., 2024). However, the ecological mechanisms underlying these observations in the tropical ocean remain poorly studied, particularly at equatorial latitudes (Chénard et al., 2019).

Efforts using ecological models have paved the way to understanding the ecological processes (selection, dispersal, and drift) structuring microbial communities (Gazulla et al., 2022; Huber et al., 2020; Junger et al., 2023; Logares et al., 2020; Vass et al., 2020). We have come to learn that prokaryotic community structures are relatively better explained by environmental selection than by dispersal or drift, while single-cell eukaryotes’ communities are more determined with the latter factors (Junger et al., 2023; Logares et al., 2020; Vass et al., 2020), mainly due to differences in organism and population sizes (Bie et al., 2012; Fodelianakis et al., 2021; Villarino et al., 2018), as well as the dormancy capacity of bacteria (Locey et al., 2020). The relative importance of these mechanisms changes with ocean depth due to differences in microbial abundances, environmental heterogeneity, and barriers to dispersal (Junger et al., 2023). However, although biotic interactions are a fundamental component of deterministic processes in ecological theory (Vellend, 2016), these ecological models rarely take them into account. As an alternative, co-occurrence network topological metrics have been used to capture the ecological characteristics of plankton communities (Fuhrman et al., 2015), to investigate community resilience (Moore et al., 2016; Solé & Montoya, 2001), as well as their potential responses to environmental changes (Chaffron et al., 2021). The application of network approaches to omics data has improved our understanding of potential ecological interactions between microbes, the role of keystone species, and the likely response of these communities to ocean environmental changes (Chaffron et al., 2021; Cram et al., 2015; Deutschmann et al., 2021; Krabberød et al., 2022; Lima-Mendez et al., 2015).

Here we compare the composition, stability, and diversity of microbial communities – both prokaryotes and eukaryotes – in two coastal marine time-series located at contrasting latitudes. We also aimed to compare the ecological processes, the environmental drivers, and the microbial co-occurrence network metrics between both sites. We determined which processes (deterministic selection vs. stochastic historical contingency and drift) temporally structure free-living (0.22-3 µm) and particle-associated (>3 µm) prokaryotes as well as small (<3 µm) and large (>3 µm) protists in each time-series. To do so, we sequenced 16S and 18S rRNA gene amplicons from DNA samples collected monthly during 41 months from April 2013 to August 2016 in two microbial observatories, one tropical site located in the Western Equatorial Atlantic (6°S), and one temperate site located in the Northwestern Mediterranean Sea (42°N). Tropical zones, which have limited variation in daylength and temperature over time, may represent a case of weak abiotic selection, while in temperate zones, abiotic selection is expected to change with seasons. As a consequence, tropical microbial communities should be more stable over time when compared to those of temperate regions, and historical and drift-like stochastic processes may arise as the main drivers structuring these communities. In this sense, we hypothesize that temperate microbial communities are primarily driven by abiotic selection caused by the seasonal fluctuation of physical variables. Meanwhile, in tropical regions where daylength and temperature remain nearly constant year-round, selection is exerted predominantly by multiple biotic factors.

## Material and methods

### Study sites and sampling design

Surface seawater (∼1 m depth) samples were collected monthly from April 2013 to August 2016 at two coastal marine observatories: the Equatorial Atlantic Microbial Observatory – EAMO (-5.99°,-35.08°) located in the western coast of the Atlantic, 30 km away from the city of Natal (Brazil); and the LTER Blanes Bay Microbial Observatory – BBMO (41.66°, 2.80°) located in the Northwestern Mediterranean Sea (Figure 1). The BBMO is a well-studied temperate oligotrophic coastal site ∼1 km offshore with little riverine or human influence (Gasol et al., 2016), while the EAMO is a newly established tropical oligotrophic site ∼3 km from the Brazilian coastline. Both coastal sites are on average 20 m deep. To our knowledge, the EAMO is the first microbial observatory located in the South Atlantic Ocean (Menezes et al., 2023), and one of the very few observatories in low-latitudes (0-10°) with available amplicon sequencing data. On the other hand, the BBMO is a long-standing microbial observatory with several publications using amplicon sequencing data (Auladell et al., 2022; Ferrera et al., 2024; Giner et al., 2019). These observatories have contrasting variation in environmental characteristics, notably temperature and daylength (Figure 1), and significant differences in most environmental variables (Figure S1).

**Figure 1.**
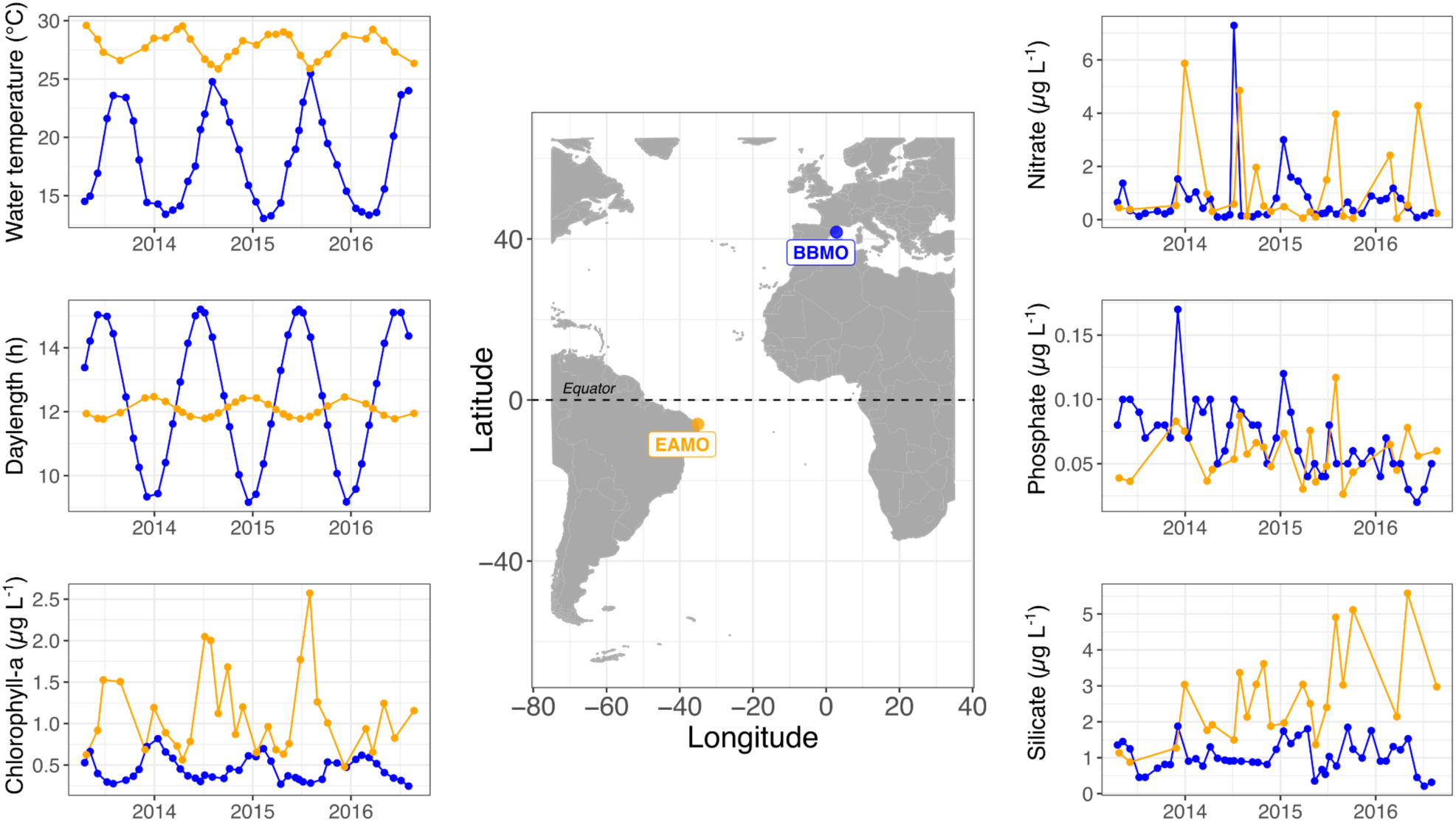
The geographic location of the two contrasting coastal marine microbial observatories sampled in this study: the Equatorial Atlantic Microbial Observatory (EAMO, 6°S) and the Blanes Bay Microbial Observatory (BBMO, 42°N). Temporal variability in temperature, daylength, chlorophyll-a, and inorganic nutrients in these observatories during the sampling period of this study (April 2013 to August 2016). Figure S1 shows the differences between observatories for all measured variables.

In each site, 20 L sub-surface (∼1 m depth) seawater samples were passed through a 200-µm mesh net and transported to the lab in 20-L in polycarbonate carboys, under dim light, and processed within 1.5 h. To obtain microbial biomass, 2 to 6 L of surface seawater was first filtered through a 20-µm filter nylon mesh and then sequentially filtered through 3 μm polycarbonate filters (hereafter “>3 μm” size-fraction; “particle-associated prokaryotes” for prokaryotes associated to particles or eukaryotic organisms, and “nanoeukaryotes” for large eukaryotes) and 0.22 μm Sterivex “cartridges” (Millipore) (hereafter “0.22-3 μm” size-fraction; “free-living” for prokaryotes and “picoeukaryotes” for small eukaryotes), using a peristaltic pump. The filters were embedded in a lysis buffer solution and maintained in the frozen (-80 °C) until DNA extraction.

Daylength (hours of light) was calculated for both sites based on coordinates and sampling dates. The accumulated rainfall data (Figure S2) was obtained from the closest meteorological stations through the National Institute of Meteorology (*INMET*, Brazil) for EAMO, and through the Catalan Meteorological service (www.meteo.cat) for the BBMO. To evaluate the potential influence of inter-annual changes on community composition in the EAMO and the BBMO, monthly values of the Southern Oscillation Index (SOI) and the Western Mediterranean Oscillation Index (WeMOI) were obtained from the NOAA database (https://www.ncei.noaa.gov/access/monitoring/enso/soi), and the University of Barcelona Climatology Group database (Lopez-Bustins et al., 2020), respectively (Figure S2).

### Analytical methods

Water transparency was determined with a Secchi disk. Water temperature and salinity were measured *in situ* with a multiparameter probe (Horiba U-50 Series) in the EAMO, and a CTD model SAIV A/S SD204 in the BBMO. *In situ* chlorophyll-a was measured at the BBMO (not measured at the EAMO) by filtering the seawater in 3 µm polycarbonate filters and then GF/F (Whatman), and extracted with acetone (90% acetone, 4°C, overnight), and determined by fluorometry (Yentsch & Menzel, 1963). Satellite-based chlorophyll-a estimates were obtained from the ESA OceanColour-CCI database (OC-CCI5; https://www.oceancolour.org/) for both sites with a daily 3×3 pixel box. Then, a 30-day moving average was calculated for each sampling date of both sites. Since satellite and *in situ* chlorophyll-a estimates were strongly correlated (Pearson’s r= 0.78, p<0.01) at the BBMO (Figure S3), the satellite-derived chlorophyll-a was used as a standard comparative proxy of chlorophyll-a between the observatories. Dissolved inorganic nutrient (NO ^−^ NO_2_^−^, NH_4_^+^, PO_4_^3−^, SiO_2_) concentrations were determined from GF/F (Whatman) filtered seawater samples in an autoanalyzer following standard procedures (Grasshoff et al., 2009).

Seawater samples for flow cytometry counts were preserved with 1% paraformaldehyde + 0.05% glutaraldehyde (final concentration). Bacterial abundance was determined in a BD FACSCalibur flow cytometer with a blue (488 nm) laser and SybrGreen I staining, according to (Gasol & del Giorgio, 2000). Picocyanobacteria were subtracted from independent counts of non-stained samples in a plot of side light scatter versus red and orange fluorescences. Flow cytometric data acquisition and analysis were performed with the FlowJo^®^ V10.0.8 software. Bacterial production rates were measured using the [^3^H]-leucine incorporation method (Kirchman, 1992). Further details on the analytical methods are provided in the Supplemental Information.

### DNA extraction, sequencing, and bioinformatics

DNA extraction was carried out with a phenol-chloroform protocol (Logares et al., 2014), and subsequent purification using Amicon columns (Millipore® 100KDa/100.000MWCO), following the manufacturer instructions. DNA extracts were quantified with a Qubit 1.0 (Thermo Fisher Scientific) and preserved at −80 °C. PCR amplification was performed using the primers 515F-Y (5’-GTGYCAGCMGCCGCGGTAA) and 926R (5’-CCGYCAATTYMTTTRAGTTT) for the 16S rRNA gene hypervariable V4-V5 region (≈ 400 bp) to target prokaryotes – both Bacteria and Archaea (Parada et al., 2016). For prokaryotes, a mock community with 25 bacterial and 2 archaeal strains (Supplemental Information, Table S1) was included as a positive control in the sequencing batch. For eukaryotes, the primers used were TAReukFWD1 (5’-CCAGCASCYGCGGTAATTCC-3’) and TAReukREV3 (5’-ACTTTCGTTCTTGATYRA-3’) of the 18S rRNA gene hypervariable V4 region (≈ 380 bp) (Stoeck et al., 2010). Negative controls (extraction and PCR blanks) were included in both sequencing batches.

Samples were sequenced in an Illumina MiSeq platform (2 x 300 bp) in two different sequencing runs (Supplemental Information). Raw reads were processed using DADA2 (Callahan et al., 2016) to determine amplicon sequence variants (ASVs). For the 16S rRNA gene, we trimmed the forward reads at 210 bp, and the reverse reads at 180 bp, while for the 18S rRNA gene, forward reads were trimmed at 220 bp and the reverse reads at 190 bp. Afterwards, for the 16S rRNA gene, the maximum number of expected errors (maxEE) was set to 5 for the forward and reverse reads, while for the 18S rRNA gene, the maxEE was set to 5 and 6 for the forward and reverse reads, respectively. Finally, error rates were estimated using DADA2 for both the 16S and 18S genes to delineate the ASVs.

ASVs taxonomy was assigned with DADA2 using the naïve Bayesian classifier method (Wang et al., 2007) together with the SILVA v.138 database (Quast et al., 2013) for prokaryotes, and the Protist Ribosomal Reference database (PR^2^, version 4.14, (Guillou et al., 2013)), for eukaryotes. Eukaryotes, chloroplasts, and mitochondria were removed from the 16S ASVs table, while Holozoan (Metazoa and Fungi), Streptophyta, and nucleomorphs were removed from the 18S ASVs table. A total of 4,922,683 reads were obtained for the 16S rRNA and 18,844,773 reads for the 18S rRNA after quality filtering and removal of the above-mentioned sequences. The samples’ prokaryotic and eukaryotic taxonomic compositions, segregated both by observatory and size-fraction, are presented in the Supplemental Information (Figure S4). For ecological models and statistical analyses, the prokaryotic and eukaryotic ASV tables were rarefied down to 8,063 and 9,513 reads per sample, respectively, using the *rrarefy* function from the *vegan* R package (Oksanen et al., 2024). For network analyses, the prokaryotic and eukaryotic read abundance tables were not rarefied but were separately subjected to a centered-log-ratio transformation before being merged to account for data compositionality (Gloor et al., 2017).

### General statistical analyses

We used the R software v 4.0.3 (R Core Team, 2013) with the packages *vegan* (Oksanen et al., 2024) and *BiodiversityR* (Kindt & Coe, 2005) for data processing and statistical analyses. In order to detect differences in community composition between sites, Bray-Curtis dissimilarity distances between sites using ASVs’ relative abundances were calculated and plotted in a non-metric multidimensional scaling plot using the *vegan* package in R. Statistical differences between sites, size-fractions, and seasons were tested by Permutational MANOVA (PERMANOVA) analysis with the ‘adonis2()’ function in the *vegan* package. Analyses of dissimilarities were also conducted using the ‘adonis2()’ function of the *vegan* R package to investigate the percentage of variance in community composition explained by biological (prokaryotic vs. eukaryotic community dissimilarity matrices) and environmental variables (McArdle & Anderson, 2001).

### Defining cosmopolitan and indicator ASVs

Common and unique ASVs were found with Venn plots using the *ggvenn* R package. We categorized the common ASVs with more than 50% occurrence in both sites as “cosmopolitan”. Since some common ASVs were significantly more abundant in a given site, we carried out a taxa indicator analysis to identify ASVs that were not necessarily unique, but were strongly associated to a given site/size-fraction (Cáceres & Legendre, 2009). We performed this indicator analysis using the ‘multipatt()’ function of the R-package *indicspecies* v.1.7.8 with parameter “IndVal.g.”, with an ASV considered as an indicator of a given site/size-fraction when the association value was >0.7, and the p-value was <0.01. The ASVs only present in the EAMO and the BBMO were called as “EA-exclusive” and “BB-exclusive”, respectively. The remaining ASVs that did not fall into any of the above categories were classified as “background”.

### Determining seasonal ASVs

Seasonality was evaluated by testing the occurrence of periodic patterns in the ASVs from both the 16S and 18S rRNA gene datasets, at each size-fraction and site. We used the Lomb Scargle periodogram (LSP) implemented in the *lomb* v.2.5.0 R-package, as it has been successfully applied in previous studies with uneven sampled signals (Auladell et al., 2022; Ferrera et al., 2024; Lambert et al., 2019; Ruf, 1999). For each dataset, we analyzed the ASVs appearing in at least 10% of samples, and applied the LSP separately using the “randlsp()” function with parameters: normalize = “standard” and type=”period”. After a manual inspection (Supplemental Information), we considered an ASV as seasonal when it presented a PN >0.2, a p-value <0.01 and an interval of ∼1 year.

### Computation of ecological processes

Phylogenetic trees were built for both the 16S and 18S rRNA gene datasets to obtain the phylogenetic distances for the ecological model (see section below). First, we aligned raw ASV sequences against an aligned SILVA template – for 16S rRNA – and an aligned PR^2^ template – for 18S rRNA – using mothur (Schloss et al., 2009). Poorly aligned regions or sequences were removed using *trimAl* (parameters:-gt 0.3-st 0.001) (Capella-Gutiérrez et al., 2009). The alignment was visually curated with *seaview* v4 (Gouy et al., 2010), and sequences with >=40% of gaps were removed. Phylogenetic trees were inferred from the curated alignment using FastTree v2.1.9 (Price et al., 2009).

We estimated the relative importance of ecological processes shaping microbial communities using a null model approach (Stegen et al., 2013) that has been widely used in ecological studies (Gazulla et al., 2022; Huber et al., 2020; Junger et al., 2023; Logares et al., 2020; Vass et al., 2020). This analysis consists of inferring environmental selection from ASV phylogenetic turnover (Table 1, Figure S5). First, we determined the phylogenetic turnover using the abundance-weighted β-mean nearest taxon distance (βMNTD) metric (Stegen et al., 2013), which computes the mean phylogenetic distances between each ASV and its closest relative in each pair of communities (pairwise comparisons). Second, we run null models with 999 randomizations to simulate the community turnover by chance (βMNTD_null_) (Stegen et al., 2013). Finally, the β-Nearest Taxon Index (βNTI) was calculated from the differences between the observed βMNTD and the mean βMNTD_null_ values. Overall, |βNTI| > 2 indicates that taxa are phylogenetically more related or less related than expected by chance, pointing to a strong influence of selection on community assembly (Stegen et al., 2013). More precisely, βNTI values higher than 2 indicate the action of heterogeneous selection, while βNTI values lower than –2 point to the action of homogeneous selection (Stegen et al., 2013). The fraction of β-diversity of the communities that was not explained by selection (|βNTI| ≤ 2) was assessed in a second step. This typically involves calculating ASV taxonomic turnover to evaluate the roles of dispersal and ecological drift in shaping community structure (Stegen et al., 2013). However, because dispersal cannot be directly inferred from temporal samples collected at a fixed site, we redefined dispersal as “non-selective turnover” (hereafter referred to as “non-selection”), dispersal limitation as “historical contingency” (Fukami, 2015), and homogenizing dispersal as “non-selective low turnover”, adapted for a temporal framework (Table 1). We computed the Raup-Crick metric (RCbray) based on Bray-Curtis dissimilarities (Chase & Myers, 2011; Stegen et al., 2013). The RCbray metric compares observed β-diversity to that derived from null models generated through 999 randomizations, representing random community assembly or ecological drift. RCbray values range from −1 to 1, with 0 indicating no deviation from the null expectation. A threshold of |RCbray| > 0.95 (two-tailed test, α = 0.05) indicates that two communities differ significantly from the null model (Stegen et al., 2013). Values of |RCbray| ≤ 0.95 suggests community assembly driven solely by ecological drift (i.e., random processes) (Chase & Myers, 2011), while RCbray > 0.95 or < −0.95 indicates that community assembly is structured by historical contingency or non-selective low turnover, respectively (Table 1).

**Table 1.**
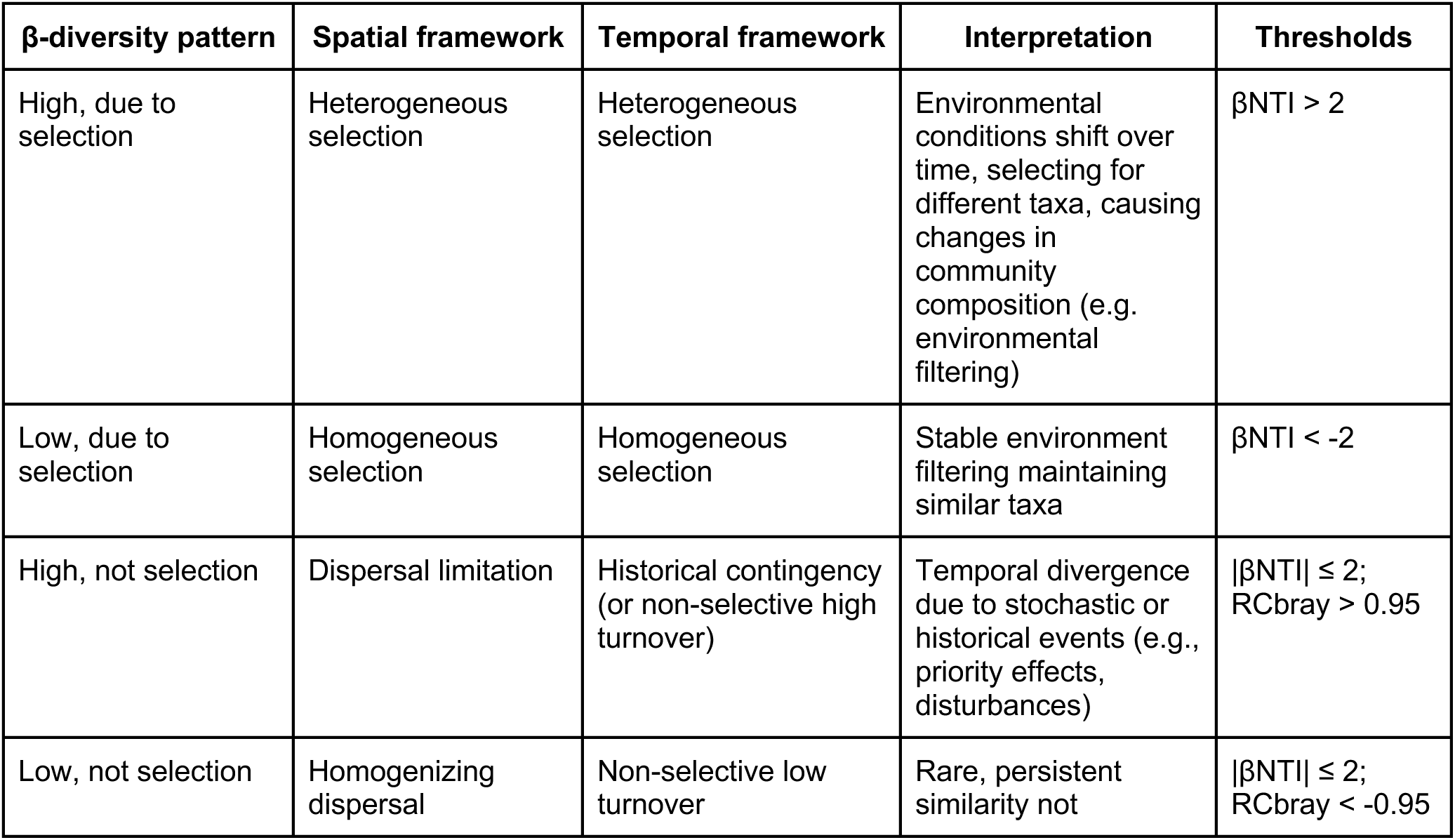
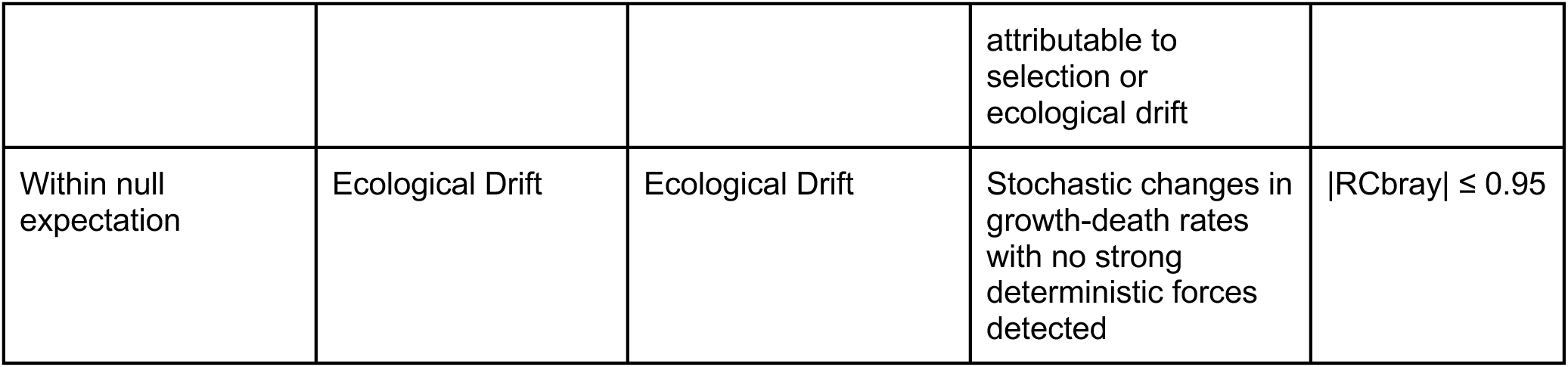
Definition of ecological processes in a temporal framework (adapted from the spatial framework in (Stegen et al., 2013). A schematic figure is available in the Supplemental Information (Figure S5).

### Network construction

Microbial association networks consist of nodes representing microorganisms and edges indicating potential interactions. For network construction, we used only samples that had both 16S and 18S data and “small” (0.22–3 µm) and “large” (>3 µm) size-fractions (Table S2, Figure S6). First, to control for data compositionality in network construction (Gloor et al., 2017), we applied a centered-log-ratio transformation separately to the prokaryotic and protist ASV tables of both size-fractions and locations. Second, we merged the four data tables corresponding to each primer/size-fraction (16S 0.22-3 µm, 16S >3 µm, 18S 0.22-3 µm and 18S >3 µm) into a single matrix for each observatory. Then, we constructed one global network for each observatory (BBMO and EAMO) using FlashWeave (Tackmann et al., 2019), selecting the options “heterogeneous” and “sensitive”. Next, we filtered environmentally driven edges via EnDED, and edges with a co-occurrence score below 50% (Deutschmann et al., 2021). Isolated nodes, i.e., nodes without an edge, were removed. The resulting (static) BBMO network contained 1906 nodes and 2614 edges (2393 or 92% positive, and 221 or 8% negative), while the (static) EAMO network contained 2074 nodes and 2389 edges (2068, 87% positive, and 321, 13% negative) (Table S2). Finally, we approximated the temporal networks via monthly subnetworks (23 in the BBMO and 22 in the EAMO) from each static network (Deutschmann et al., 2023). Each temporal network contains a subset of nodes and edges of the global static network. An edge is present in a subnetwork of a particular month if it was present in the static network and both nodes correspond to microorganisms detected in that month. A microorganism is determined as detected if the sequence abundance is above zero.

### Network analysis

We computed global network metrics to characterize the single static network and each monthly subnetwork using the *igraph* R-package (Csardi & Nepusz, 2006) and adapted code from (Deutschmann et al., 2023, 2024). The computed metrics included edge density, average path length, transitivity, mean degree, and assortativity based on node degree, domain (prokaryotes vs. protists), and size-fraction (small vs. large). We also calculated the average strength of positive associations between nodes. The definitions and ecological interpretations of these metrics are summarized in Table 2. Spearman correlations between global network metrics and environmental data were computed using the *corr.test* function in the *psych* R package (Revelle, 2025), with Holm’s correction for multiple testing (Holm, 1979). Network visualizations were generated with Gephi v.0.10.1 (Bastian et al., 2009) using the Fruchterman–Reingold layout (Area = 10,000; Gravity = 1; Speed = 10).

**Table 2.**
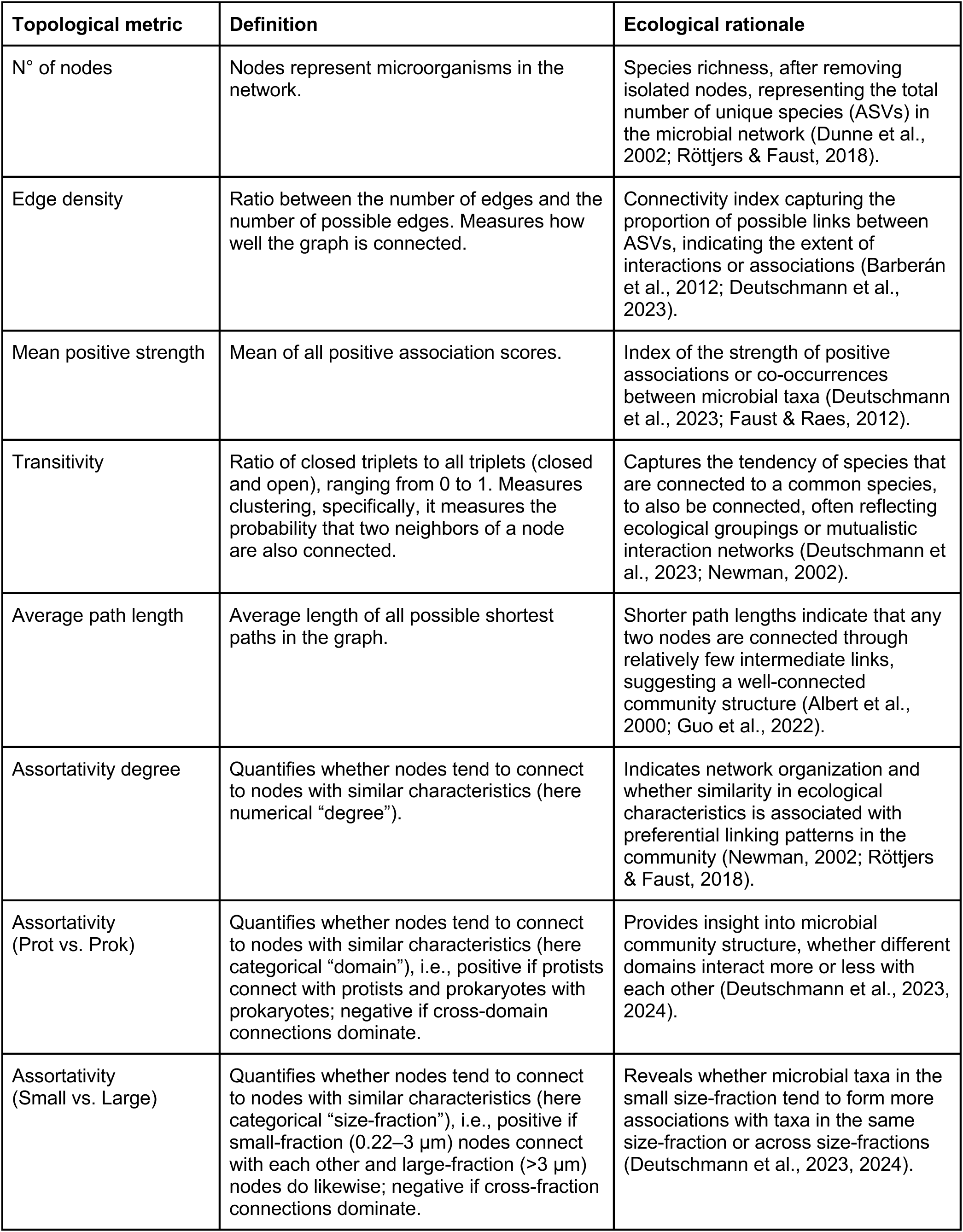
Definition and ecological rationale of the network topological metrics computed in this study.

## Results

### Equatorial microbial communities exhibit lower turnover than those at the temperate observatory

Alpha diversity analyses revealed significantly lower prokaryotic diversity at the tropical site relative to the temperate site (Mann-Whitney Rank Sum test, p<0.01; Figure 2A and S7). Significant differences in compositional diversity were recorded between prokaryotic size-fractions (ANOVA, p<0.001). In contrast, small protist diversity was greater at the EAMO than at the BBMO (Mann-Whitney test, p<0.001), although this pattern did not extend to larger protists (Figure 2A and S7). Seasonal variation in alpha diversity was evident at the BBMO but not at the EAMO (Figure S8). Prokaryotic diversity peaked during winter, while protist diversity was highest in autumn at the BBMO. Conversely, the EAMO displayed no significant seasonal fluctuations in alpha diversity, consistent with a more temporally stable tropical microbiome. Rank-abundance curves further illustrated these differences: the BBMO communities were dominated by fewer, highly abundant ASVs, whereas the EAMO harbored a greater number of less abundant taxa, a pattern especially pronounced for protists (Figure 2B).

**Figure 2.**
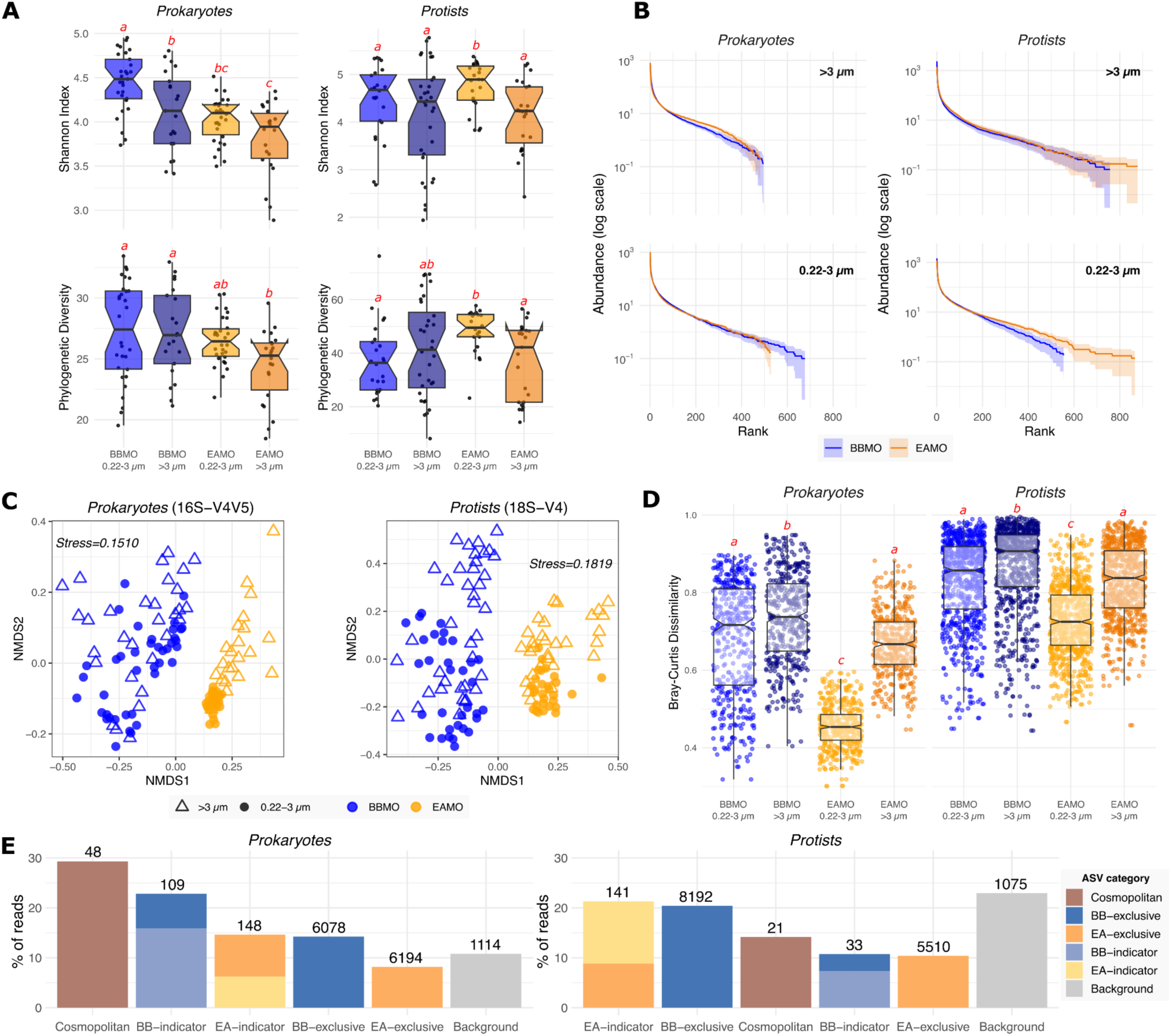
Differences in microbial community diversity between the temperate site (BBMO) and the tropical site (EAMO). **A)** Shannon index and phylogenetic diversity (Figure S7 includes ASV richness, and Pielou’s evenness, while Figure S8 shows seasonal variability in diversity metrics). Different red letters represent significantly different means (ANOVA, Tukey post hoc test, *p* < 0.05); **B)** Normalized rank abundance plots of the prokaryotic and protist communities. The line stands for the average rank abundance of each fraction and site. The area around the lines represents rank abundance dispersion of each sample subset; **C)** Nonmetric multidimensional scaling (NMDS) based on the Bray-Curtis dissimilarities among prokaryotic and eukaryotic samples – labeled by observatory (colors) and size-fraction (shapes); **D)** Beta-diversity between observatories and size-fractions; **E)** Percentage of reads of the ASV categories, as defined in the methods section. The number (#) of ASVs within each category is given above the bars. Note that some exclusive ASVs were also indicators for prokaryotes (n = 51 in BB; n = 91 in EA) and protists (n = 19 in BB; n = 80 in EA). BB = BBMO, Blanes Bay Microbial Observatory; EA = EAMO, Equatorial Atlantic Microbial Observatory.

Beta diversity analyses showed strong segregation of microbial community composition by site (Figure 2C), which was a more informative factor than size-fraction in explaining variability for both prokaryotic (PERMANOVA, site *R*^2^ = 0.22, size-fraction *R*^2^ = 0.06, *p* < 0.001) and protist (site *R*^2^ = 0.12, size-fraction *R*^2^ = 0.05, *p* < 0.001) communities. The equatorial site exhibited significantly lower temporal turnover (Bray-Curtis dissimilarity) in both domains and size-fractions compared to the temperate site (Figure 2D). At BBMO, seasonality explained more variation than size-fraction for both prokaryotic (PERMANOVA, season *R*^2^ = 0.29, size-fraction *R*^2^ = 0.07, *p* < 0.001) and protist (season *R*^2^ = 0.14, size-fraction *R*^2^ = 0.07, *p* < 0.001) communities (Figure S9). In contrast, at EAMO, seasonality accounted for less variation than size-fraction in the prokaryotic community (season *R*^2^ = 0.09, size-fraction *R*^2^ = 0.19, *p* < 0.001), whereas in the protist community, seasonality (*R*^2^ = 0.11) and size-fraction (*R*^2^ = 0.08, *p* < 0.001) explained a similar fraction of the variation.

### Higher stability and number of microbial indicators in the equatorial site

Despite a similar percentage of shared ASVs between biological domains and size-fractions (∼8–9%; Figure S10), marked differences emerged in the distribution and identity of cosmopolitan and site-specific indicator ASVs. Among shared particle-associated prokaryotic ASVs (n = 788), 23 (∼3%) were classified as cosmopolitan, while among free-living prokaryotic ASVs (n = 703), 41 (∼6%) were cosmopolitan, with 16 shared across both size-fractions, 25 exclusive to the free-living fraction, and 7 to the particle-associated fraction. For protists, cosmopolitans were rarer: 6 (∼0.8%) of 768 ASVs in the large size-fraction and 16 (∼2%) of 832 ASVs in the small size-fraction, with only two ASVs found to be cosmopolitan in both size-fractions. The percentage of unique prokaryotic ASVs in each observatory was similar between size-fractions, with ∼46% in the BBMO, and ∼45% in the EAMO (Figure S10). In contrast, the BBMO harbored more unique protist ASVs than the EAMO in both fractions (small: 50% vs. 41%; large: 53% vs. 39%; Figure S10–12). The indicator species analysis revealed more indicator ASVs at EAMO (148 prokaryotes, 141 protists) than BBMO (109 and 33, respectively) (Figure 2E). Many site-exclusive ASVs were also indicators: at EAMO, 91 prokaryotes (∼61%) and 80 protists (∼57%); at BBMO, 51 prokaryotes (∼47%) and 19 protists (∼58%).

The 30 most abundant prokaryotic ASVs highlighted contrasting temporal dynamics (Figure 3). At BBMO, strong seasonality and niche partitioning were evident for cosmopolitan and indicator taxa. For example, *Synechococcus* CC9902 cosmopolitan ASVs peaked in summer, while BBMO-specific ASVs dominated in early spring; Flavobacteriaceae ASVs (e.g., NS4/NS5 groups, *Formosa*) increased in autumn/winter, and ASVs assigned to taxa such as *Balneola*, *Aurantivirga*, and *Erythrobacter* were largely confined to the BBMO. In contrast, EAMO communities were more temporally stable, with cosmopolitan (*Synechococcus* CC9902, Flavobacteriaceae, SAR11, *Cyanobium* PCC-6307) and site-indicator (*Actinomarinales*, SAR406, *Cyanobium* PCC-6307) ASVs showing little temporal change. An exception was a cosmopolitan ASV assigned to the genus *Fluviicola*, which exhibited more striking temporal variability in the larger size-fraction at EAMO. Notably, ASV-level niche partitioning was observed in *Synechococcus* CC9902, with some ASVs acting as generalists and others showing site-specific dominance.

**Figure 3.**
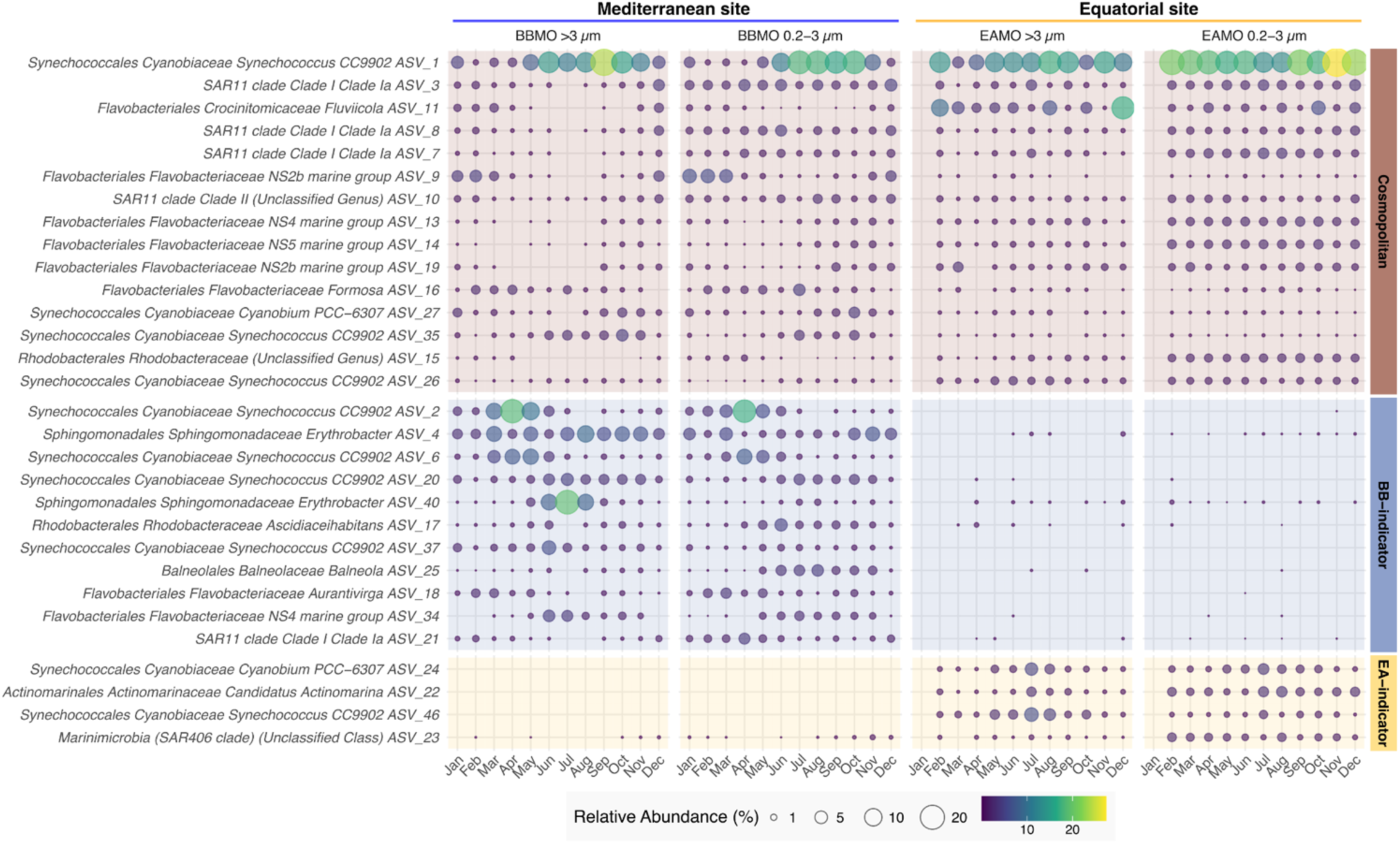
The monthly average relative abundance of the 30 most abundant prokaryotic ASVs classified as cosmopolitan, BB-indicators, or EA-indicators. BB = BBMO, EA = EAMO. A version of this figure with the 100 most abundant prokaryotic ASVs is presented in the Supplemental Information (Figure S13).

The differences in seasonal and niche-partitioning patterns between sites were even clearer in protists (Figure 4). At BBMO, distinct seasonal patterns and niche partitioning appeared among both cosmopolitan and indicator protist ASVs. For example, ASVs assigned to the cosmopolitan *Bathycoccus prasinos* and the BBMO-indicator *Micromonas bravo* B1 peaked in winter and early spring, while other cosmopolitan *Micromonas* clades (*M. bravo* B2 and *M.* clade B4) slightly increased during spring. The small centric diatom *Minidiscus comicus* was also more abundant in winter. Conversely, ASVs assigned to *Gyrodinium dominans* and background dinoflagellates like *Ansanella granifera* and *Heterocapsa* peaked in warmer months, likely associated with seasonal stratification and low nutrient availability. The cryptophyte *Teleaulax gracilis* had moderate abundance peaks in spring, while BBMO-indicator ASVs assigned to the cryptophyte *Plagioselmis prolonga* and green algae *Chlorodendrales* increased in summer. Parasitic dinoflagellate (Syndiniales) ASVs from Dino-Group-I-Clade-1 showed temporal niche partitioning, with early and late spring peaks for different ASVs, whereas Dino-Group-I-Clade-4 increased in autumn.

**Figure 4.**
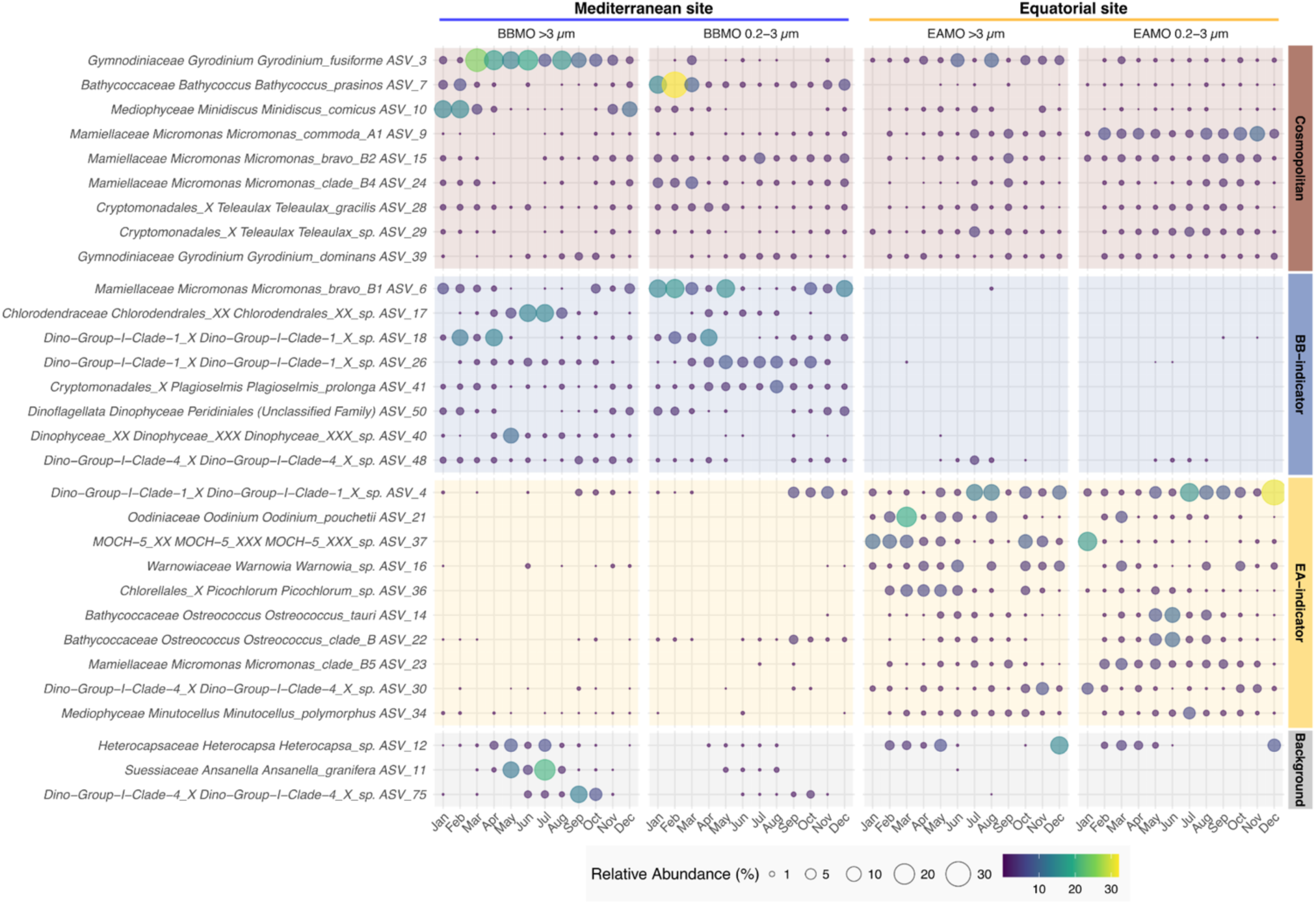
The monthly average relative abundance of the 30 most abundant protist ASVs classified as cosmopolitan, BB-indicators, EA-indicators, or background. BB = BBMO, EA = EAMO. A version of this figure with the 100 most abundant protist ASVs is presented in the Supplemental Information (Figure S14).

Although less pronounced than at BBMO, protist community temporal fluctuations were still observed at the EAMO (Figure 4). Cosmopolitan *Micromonas* clades (*M. commoda* A1, *M. bravo* B2, and *M.* clade *B4*) as well as the EAMO-indicator *Micromonas* clade B5, were present year-round with moderate abundance and minimal fluctuation. In contrast, EAMO-indicators *Ostreococcus tauri* and *Ostreococcus* clade B peaked mid-year, coinciding with the region’s rainy season (Figure S2). Cosmopolitan *Gyrodinium fusiforme* and *G. dominans* persisted across seasons with little variation. The small diatom *Minutocellus polymorphus* was consistently detected year-round, with a mid-year peak. Syndiniales parasites belonging to Dino-Group-I-Clade-1 and - Clade-4, along with *Oodinium pouchetii* and the athecate dinoflagellate genus *Warnowia*, were also persistent throughout the year. Among EAMO-indicators, ASVs assigned to the phototrophic Marine Ochrophyta clade 5 (MOCH-5) and the green algae *Picochlorum* showed some temporal variation, with relative abundances dropping mid-year.

### Weaker seasonal patterns in the equatorial site than in the temperate site

Our LSP (Lomb Scargle periodogram) analysis to detect seasonal ASVs revealed pronounced differences in their temporal dynamics between the two sites. From the total of 180 ASVs identified as seasonal across all datasets, the vast majority (85%, or 153 ASVs) belonged to the temperate site BBMO, while only 27 seasonal ASVs (15%) were detected at the equatorial site EAMO (Figure 5A). The number of seasonal ASVs was comparable across domains (96 prokaryotes vs. 84 protists) and across size-fractions, indicating that the pronounced seasonal signal in the BBMO was not limited to one group or size-fraction. At the BBMO, seasonal prokaryotes accounted for ∼9% of the total reads (6.54% in the free-living and 11.38% in the particle-associated fractions), while in the EAMO they represented <3% of the reads (4.63% and 0.15%, respectively). Similarly, seasonal protists comprised 8.12% of the total reads at the BBMO (9.63% for small protists and 6.6% for large protists), compared to just 3.27% in the EAMO. Strikingly, none of the large protist ASVs at the EAMO passed the seasonality threshold in a sharp contrast with the BBMO.

**Figure 5.**
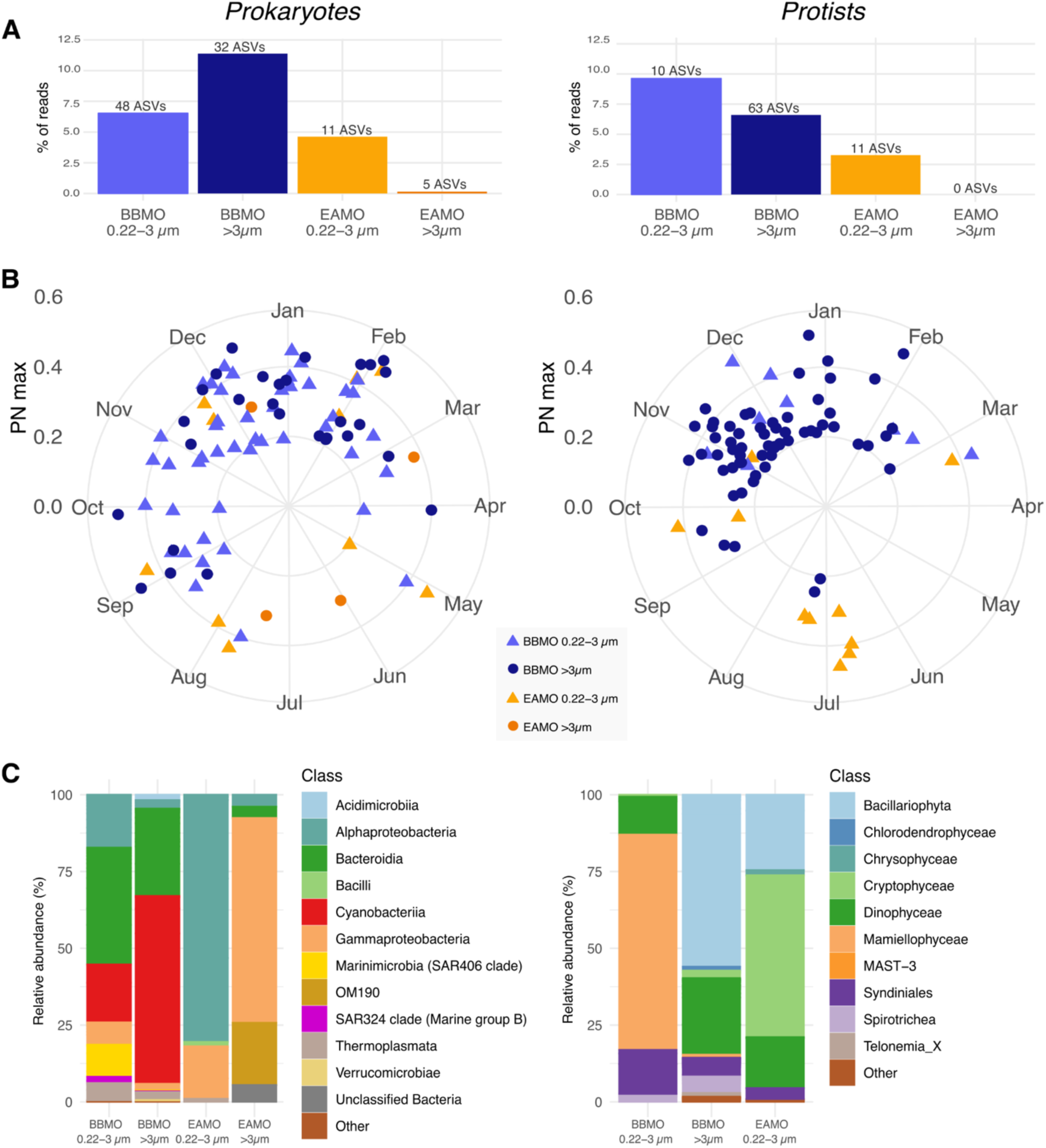
Seasonal ASVs in the temperate site (BBMO) and the tropical site (EAMO). **A)** Percentage of reads and number of seasonal ASVs in each site and size-fraction. **B)** Polar plots representing the seasonal ASVs of the prokaryotic and protist communities. Higher strength recurrence values (PN max) represent stronger seasonal signals, and the polar plot indicates the month in the year when the maximal abundance occurs. Figure S15 shows the same, but with seasonal ASVs coloured by their taxonomic groups. **C)** Taxonomy of the seasonal ASVs for prokaryotes (left) and protists (right) at each site and size-fraction. Note that not a single ASV could be considered seasonal in the EAMO protist large size-fraction. BBMO = Blanes Bay Microbial Observatory; EAMO = Equatorial Atlantic Microbial Observatory.

For each seasonal ASV, we extracted the month of the year with maximal abundance (Figure 5B). Seasonal ASVs at the BBMO displayed strong winter dominance, with their peak abundances occurring mainly between September and March—particularly in December, January, and February. Most seasonal prokaryotes in the BBMO were associated with the free-living fraction and included taxa from Alphaproteobacteria, Bacteroidia, and Gammaproteobacteria (Figure 5C). Interestingly, only one *Synechococcus* CC9902 ASV (Cyanobacteria) was detected as seasonal in the free-living fraction (peaking in April), while three others were seasonal in the particle-associated fraction. Groups such as Thermoplasmata (6 ASVs) and SAR324 clade (3 ASVs) exhibited clear seasonal peaks in the winter months, whereas other ASVs showed more variable timing (Figure S15). For protists, most seasonal ASVs at the BBMO were in the large fraction (>3 µm) taxa that peaked in late autumn and winter (November, December, and January). These included several diatoms (Bacillariophyta), dinoflagellates (Dinophyceae), and parasitic taxa such as Syndiniales (Figure 5C). Among the few small protist seasonal ASVs, *Bathycoccus prasinos* (ASV_7, Mamiellophyceae) stood out, with winter peaks in February that accounted for up to 30% of the total protist reads in some samples.

In contrast, seasonal ASVs in the EAMO showed no clear temporal clustering (Figure 5B). Prokaryotic seasonal ASVs were scattered throughout the year, with individual ASVs peaking in different months. For example, SAR11 (ASV_8) peaked in December, Haliaceae (ASV_600) in July, and a Rhodobacteraceae ASV (ASV_4640) in March. Interestingly, among the few seasonal protists in the EAMO, half of the small size-fraction ASVs (6 out of 11) peaked in July (rainy period), and were primarily members of Cryptophyceae, Bacillariophyta, and Dinophyceae (Figure 5C). Still, these patterns were less coherent than those observed at the BBMO, highlighting the absence of strong seasonal structuring in the equatorial community.

### Equatorial community variation is driven more by biological and stochastic processes than by seasonal selection

We observed striking differences in the relative importance of ecological processes structuring microbial communities between the two observatories (Figure 6A). Across all comparisons, deterministic selection explained a higher proportion of community turnover in prokaryotes than in protists, regardless of site or size-fraction. Moreover, selection consistently played a stronger role in the free-living (0.22–3 µm) fraction than in the particle-associated (>3 µm) one. Selection accounted for ∼26% of prokaryotic turnover in the particle-associated fraction, and ∼35% in the free-living fraction at the BBMO. Although overall selection was lower at the EAMO, the same pattern held: it explained ∼5% of turnover in the particle-associated fraction and ∼4% in the free-living fraction. For protists, selection explained a smaller fraction of the variation in both the BBMO (∼6% in both size-fractions), and the EAMO (8% in the larger and 2% in the smaller size-fraction).

**Figure 6.**
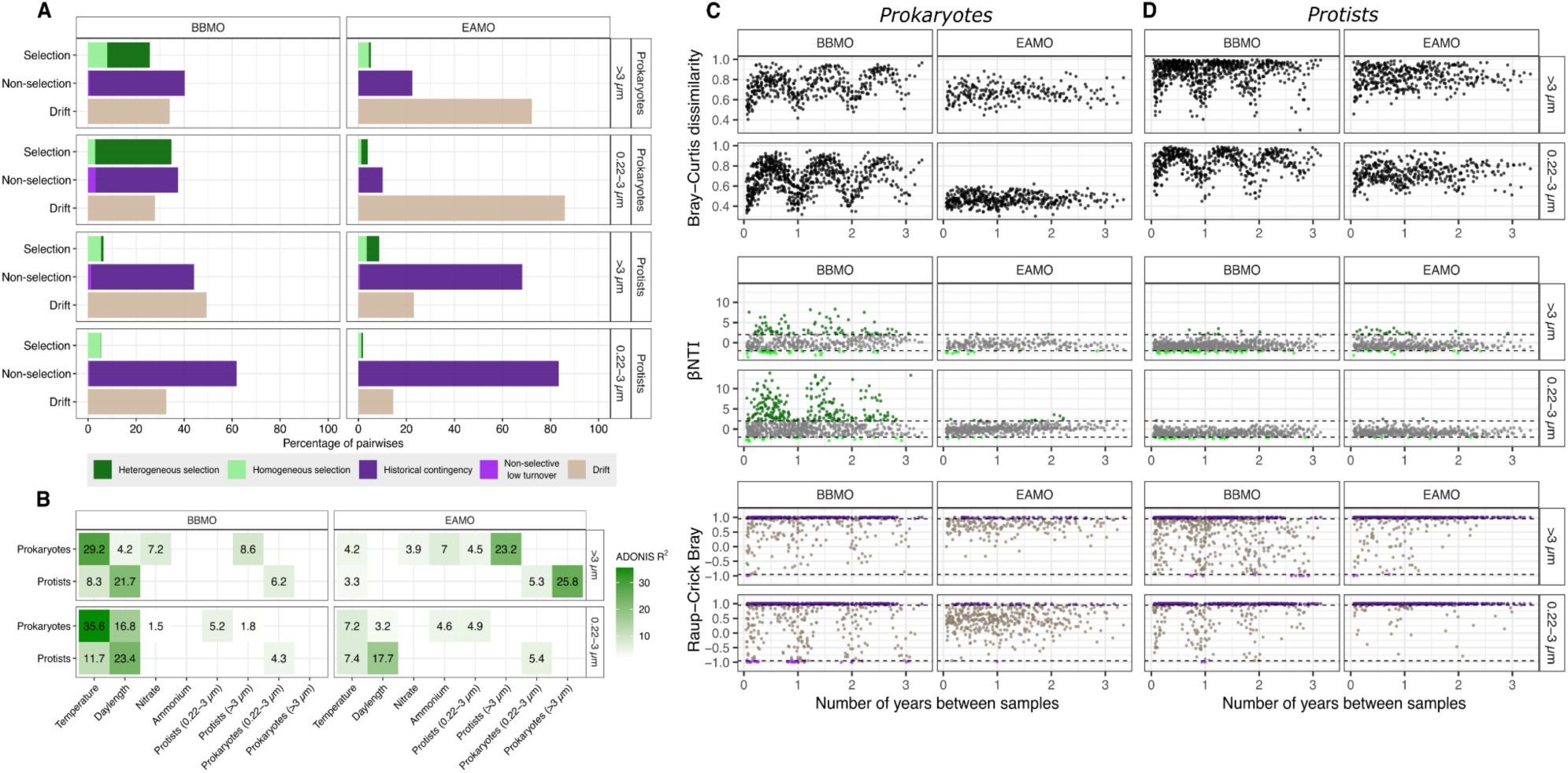
Microbial community assembly processes and environmental drivers in the contrasting time-series. **A)** Relative importance of the different ecological mechanisms structuring the microbial communities in the BBMO and the EAMO. **B)** Percentage of variance (Adonis R^2^) in eukaryotic and prokaryotic community composition (Bray-Curtis dissimilarity) explained by the biological and environmental variables with significant values (p<0.01) in at least one of the subsets. Blank spaces depict non-significant results (p>0.01). **(C, D)** Time-decay in prokaryotic and protist communities in the temperate and tropical time-series. Bray-Curtis dissimilarity, βNTI and Raup-Crick Bray between samples for each size-fraction of **C)** prokaryotes and **D)** protists plotted against the time lag between samples. BBMO – Blanes Bay Microbial Observatory. EAMO – Equatorial Atlantic Microbial Observatory.

To investigate the biological and environmental factors determining the structure of the microbial communities in the contrasting latitudes, we used dissimilarity analysis (ADONIS). The temperate site free-living prokaryotic community turnover was mainly explained by temperature (∼39%), daylength (∼17%), and secondarily by the small protist community structure (∼5%) (Figure 6B). Conversely, in the equatorial site, the free-living prokaryotic community turnover was weakly explained by temperature (∼7%), daylength (∼3%), ammonium concentration (∼5%) and small protists (∼5%) (Figure 6B). The particle-associated prokaryotic community turnover was explained by temperature (∼29%), nitrate (∼7%), daylength (∼4%), and large protists (∼9%) in the temperate site while, in comparison, the tropical particle-associated prokaryotic community was mainly explained by the large protists (∼23%), and secondarily by the small protist community structure (5%), as well as environmental variables such as temperature (∼4%), ammonium (7%), and nitrate (∼4%) (Figure 6B). For small protists (0.22–3 µm), the main drivers at the BBMO were daylength (23%) and temperature (12%), followed by the prokaryotic community (4%). In EAMO, these values were consistently lower, with daylength explaining 17%, temperature 7%, and prokaryotes 5%. For larger protists (>3 µm), turnover at the BBMO was also largely explained by daylength (22%) and temperature (8%), whereas at EAMO, the main explanatory variable was particle-associated prokaryotes (26%), followed by free-living prokaryotes (5%) and only a minor role for temperature (3%), suggesting abiotic drivers had a negligible role in shaping large protist turnover in the equatorial site.

### Equatorial network metrics show no association with seasonal environmental factors

Microbial co-occurrence networks revealed fundamental structural differences between the equatorial and temperate sites, both in terms of composition and topological properties (Figure 7). At EAMO, the network was strongly dominated by protists (77% vs 24% prokaryotes), particularly in the small size-fraction (Figure 7A). In contrast, BBMO showed a more balanced composition (53% vs 48%), with the small fraction especially enriched in prokaryotes. Several network metrics also differed significantly between the two sites (Figure 7B). The temperate network exhibited higher edge density (t = 3.3, df = 32.09, p < 0.01), transitivity (t = 2.26, df = 32.95, p < 0.05), and domain-based assortativity (t = 4.45, df = 42.63, p < 0.01), indicating greater interconnectedness and modular structure. In contrast, the equatorial network had significantly higher mean positive association strength (t = –12.26, df = 40.11, p < 0.001) and degree-based assortativity (t = –2.48, df = 41.14, p < 0.05), suggesting that connections are concentrated among a smaller core of highly connected taxa, resulting in stronger positive associations but lower overall network complexity and modularity.

**Figure 7.**
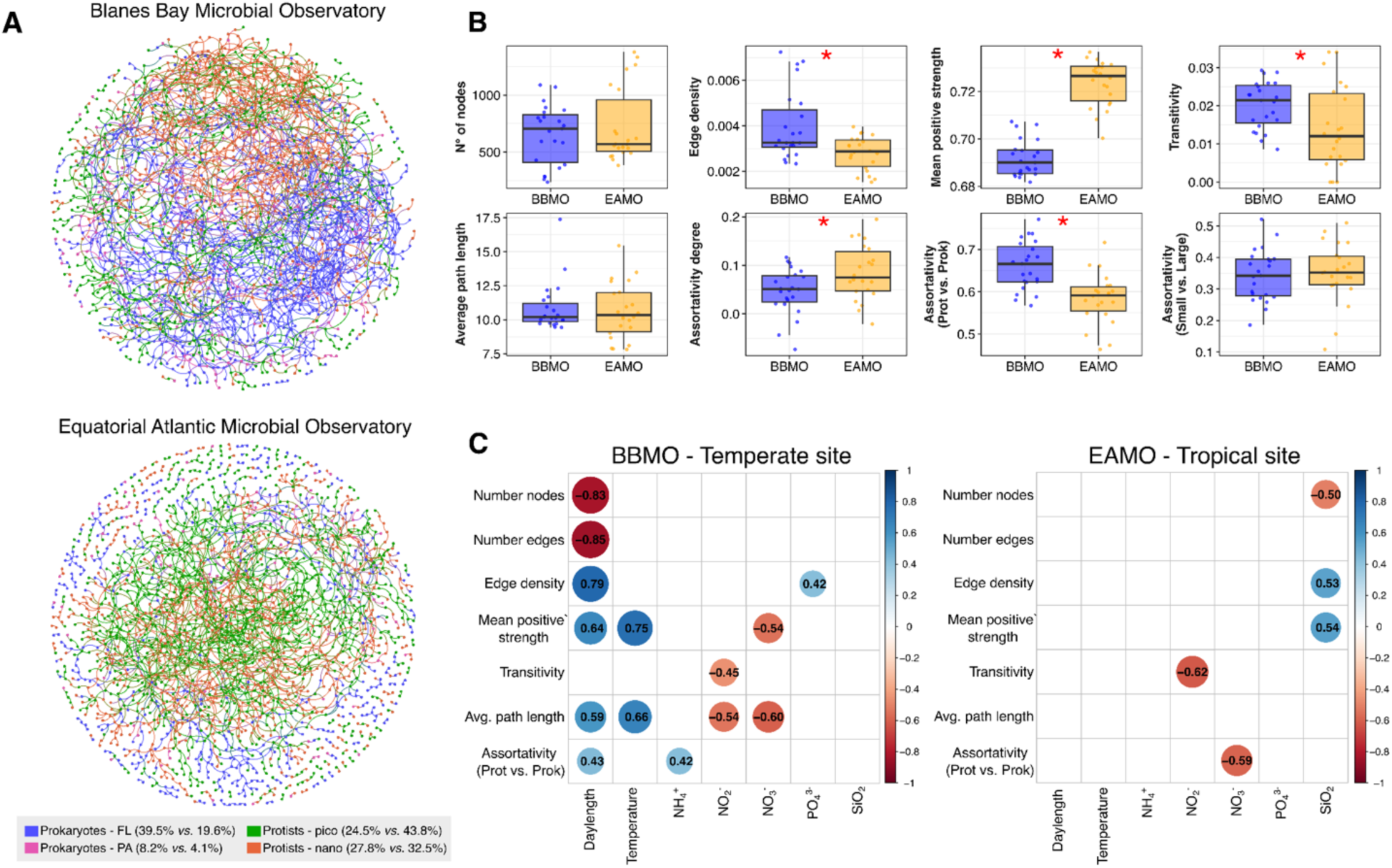
A) Visualization of the global static microbial networks of the BBMO and the EAMO. The percentages in the edge color legend indicate the proportion of the biological domain (prokaryotes and protists) and size-fraction (0.22–3 µm: free-living (FL) or picoplankton; and >3 µm: particle-associated (PA) or nanoplankton) in each network (BBMO *vs.* EAMO). Figure S16 shows the static network metrics in both observatories. **B)** The dynamic temporal network metrics differ between the BBMO and the EAMO. Red asterisks depict statistically significant differences (t-test, p<0.01). **C)** Spearman correlation matrices between network metrics and environmental variables in both observatories. Empty boxes represent non-significant correlations (p>0.05). BBMO – Blanes Bay Microbial Observatory; EAMO – Equatorial Atlantic Microbial Observatory.

A key distinction between sites was the correlation between network structure and environmental variability (Figure 7C). At BBMO, most network metrics (except transitivity) were significantly linked to seasonal factors, especially daylength and temperature. For instance, the number of nodes and edges were negatively correlated with daylength (Spearman r ≈ –0.8), while edge density and mean positive association strength were positively correlated with it (r ≈ 0.8). In contrast, EAMO showed no significant relationships with temperature or daylength, though a few metrics correlated with nutrients, such as a positive association between edge density and silicate (r = 0.53) and a negative association between transitivity and nitrite (r = – 0.62). These differences reinforce the idea that microbial community structure at low latitudes is less shaped by seasonal environmental variability.

## Discussion

Our results support our central hypothesis that microbial community turnover is less strongly governed by environmental selection in the equatorial site (EAMO) than in the temperate site (BBMO), in line with the broader expectation that environmental heterogeneity drives deterministic processes. This finding is consistent with recent reports demonstrating that selection increases with environmental variability across oceanic gradients (Junger et al., 2023; Milke et al., 2022). Comparable patterns have also been observed in microbial communities from terrestrial (Dini-Andreote et al., 2015; Stegen et al., 2013) and freshwater ecosystems (Huber et al., 2020; Vass et al., 2020). In our study, the stronger role of selection in the temperate site was supported by multiple lines of evidence: (i) a higher proportion of turnover explained by selection in the probabilistic ecological model, (ii) the greater proportion of beta-diversity explained by environmental variables such as daylength and temperature, (iii) the higher number and relative abundance of seasonal ASVs, and (iv) stronger coupling between network metrics and seasonal variables in the temperate site than in the equatorial site. Accordingly, classic studies have long noted that tropical plankton communities exhibit lower amplitude in population fluctuations, with little evidence for ecological succession or phenology (Connell & Orias, 1964; Dunbar, 1960), in contrast to the strong seasonal successions characteristic of temperate systems (Margalef, 1978). This latitudinal gradient in environmental variability likely underpins the stronger deterministic structuring and seasonal recurrence observed in the BBMO when compared to the EAMO. Additionally, higher metabolic rates in tropical environments (Amado et al., 2013) can amplify stochasticity by increasing mutation rates, intrinsic mortality, and demographic fluctuations, thereby enhancing the role of ecological drift in shaping community composition (Saito et al., 2021).

We also found marked differences between biological domains. Prokaryotic turnover was more strongly explained by selection than protist turnover across both sites and size-fractions. This aligns with previous findings from both marine (Junger et al., 2023; Logares et al., 2020) and freshwater systems (Logares et al., 2018; Vass et al., 2020), where selection more consistently structures prokaryotic communities. These differences likely stem from domain-specific traits: protists tend to be larger, with smaller population sizes, which limits their dispersal (Massana & Logares, 2013; Villarino et al., 2018) and increase the potential effects of ecological drift (Bie et al., 2012; Fodelianakis et al., 2021). Prokaryotes, by contrast, exhibit wider dispersal and a higher potential for dormancy – a mechanism less common among marine protists – which may buffer them from stochastic processes and reinforce the signature of homogeneous selection (Lennon et al., 2021; Locey et al., 2020). Nonetheless, some protists can form long-lived resistant cysts or spores, with known persistence ranging from months in the water column (Ellegaard & Ribeiro, 2018) to centuries or millennia in sediments (Sanyal et al., 2022). We also observed that selection played a relatively larger role in structuring free-living (0.22–3 µm) than particle-associated (> 3 µm) prokaryotic communities. Larger particles or hosts can provide microhabitats that buffer environmental variability (Yung et al., 2016), reducing the strength of bulk environmental selection (Mestre et al., 2020), and their spatial confinement likely imposes stronger dispersal limitation and drift on associated microbes. For instance, copepod-associated microbial communities are more shaped by drift than environmental selection than surrounding seawater communities (Velasquez et al., 2025). Moreover, we found stronger historical contingency for protists than prokaryotes in both sites. The higher number of cosmopolitan prokaryotes (n=48) compared to protists (n=21) further supports that historical contingency, mediated by dispersal limitation, is stronger for protists. Dispersal limitation strengthens historical contingency by making communities more dependent on colonization history and associated priority effects (Fukami, 2015; Nemergut et al., 2013).

Beyond ecological processes, we also observed clear differences in diversity and community composition between observatories. Prokaryotic diversity was on average significantly higher for both size-fractions at the temperate site (BBMO), particularly in winter, while protist diversity was higher only in the small size-fraction at the equatorial site (EAMO). These results partially corroborate global spatial observations showing prokaryotic diversity plateauing at mid-latitudes, and microeukaryotic diversity increasing toward the tropics (Ibarbalz et al., 2019). Time-series comparisons further indicate that the Australian temperate observatories harbor comparable or higher prokaryotic diversity than tropical sites (Raes et al., 2024), consistent with the broad optimal thermal range for prokaryotes (15–30 °C) observed at mid-to low-latitudes (Ibarbalz et al., 2019). One possible explanation for the higher mean prokaryotic diversity at the temperate site, which peaks in winter, is the stronger seasonal turnover characteristic of mid-latitudes (Fuhrman et al., 2015; Ladau et al., 2013). To our knowledge, no global direct comparisons of protist diversity across observatories have been published, and our study provides the first time-series comparison between sites at contrasting latitudes. Across both sites, protists exhibited higher temporal turnover (beta-diversity) than prokaryotes, consistent with long-term observations at the San-Pedro Ocean Time-Series (Yeh & Fuhrman, 2022).

Latitudinal differences were also evident in the number of site-indicator and seasonal ASVs. The higher number of EAMO-indicators compared to BBMO-indicators is likely explained by the greater stability in diversity at the equatorial site. Among the most abundant prokaryotic EAMO-indicators, we found ASVs assigned to *Synechococcus* CC9902, SAR406, *Cyanobium* PCC-6307, and the candidate actinobacterial genus *Actinomarina*. These bacterial genera have been previously reported in equatorial coastal waters of Singapore (Chénard et al., 2019; Wijaya et al., 2023), supporting their widespread distribution in low-latitude coastal ecosystems. We detected far fewer seasonal ASVs in the equatorial site (n = 16) than in the temperate site (n = 80). Notably, many cosmopolitan prokaryotes such as *Synechococcus* CC9902 (e.g., ASV_1 and ASV_35), *Cyanobium* PCC-6307 (ASV_27), and several Flavobacteriales (e.g., ASV_13, ASV_14, ASV_19) were seasonal in the BBMO but not in the EAMO. Conversely, only a few cosmopolitans, such as SAR11 Clade I (ASV_8) and Clade II (ASV_10), and the SAR86 clade (Gammaproteobacteria, ASV_103), along with abundant EAMO-indicators such as SAR116 (ASV_31) and OM182 (Gammaproteobacteria, ASV_116), displayed recurrent patterns at the EAMO. This reinforces the reduced influence of seasonal environmental selection in tropical waters.

Protist EAMO-indicators included several green algae assigned to Mamiellophyceae (*Micromonas* clade B5, *Ostreococcus* tauri and *O.* clade B), as well as the fast-growing, thermo-halotolerant *Picochlorum*, all found year-round in equatorial waters (Chénard et al., 2019). The aggregate-forming diatom *Minutocellus polymorphus* (ASV_79), which showed recurrent temporal patterns in the EAMO, is typically associated with coastal and estuarine environments, with documented occurrences in tropical Eastern Pacific and Indian Ocean waters (Barry-Martinet et al., 2025; Hernández-Márquez et al., 2023). The MOCH-5 group, detected here as an abundant and exclusive EAMO-indicator, has also been reported as a major phototrophic component in the tropical North Atlantic around the Amazon River plume (Charvet et al., 2021). Parasitic Syndiniales (Dino-Group-I) Clade-1 and Clade-4, both previously observed in equatorial waters (Chénard et al., 2019), had ASVs emerging as EAMO-indicators and BBMO-indicators, consistent with segregation between Mediterranean and tropical/subtropical communities (Rizos et al., 2023). Other notable EAMO-indicators showing recurrent patterns included *Oodinium pouchetii*, a parasitic dinoflagellate infecting appendicularians off the Brazilian coast (Gómez & Skovgaard, 2015), the athecate dinoflagellate genus *Warnowia*, characterized by its sophisticated photoreceptor organelle (Cooney et al., 2023), and the cosmopolitan cryptophyte *Teleaulax sp.* (ASV_29). Collectively, these abundant equatorial indicators may serve as potential candidates to monitor environmental changes (e.g., ocean warming, marine heatwaves) in temperate sites (Brown et al., 2024) such as the BBMO.

Furthermore, differences in diversity and community structure may also reflect broader ecosystem characteristics such as trophic state and continental shelf extent. The EAMO is a more productive system, with chlorophyll-a concentrations about twice those observed at the BBMO (Figure S1), where winter–spring blooms can reach ∼1 mg m^-3^ (Auladell et al., 2022). Additionally, although narrower than some global shelves, the continental shelf off the northeastern Brazilian coast where the EAMO is located is broader, averaging ∼30–40 km in width (Vital et al., 2010), than the immediate Blanes Bay coastal shelf, which is only ∼4 km wide (Durán et al., 2013). Such geomorphological differences likely enhance benthic–pelagic coupling and nutrient inputs via coastal mixing and sediment resuspension (Bauer et al., 2013), potentially promoting irregular non-seasonal variability in microbial communities at the EAMO. Indeed, studies in equatorial coastal waters have shown that monsoon seasons (Chénard et al., 2019), and local features like riverine input and tidal mixing (Wijaya et al., 2023) can drive environmental selection on microbial communities over very short time scales. In this study, the equatorial community structure was not explained by climatic variables (e.g., SOI and rainfall), and our monthly data suggest non-seasonal nutrient variation in the EAMO, but capturing these dynamics more precisely would require higher temporal resolution than in our study. Future studies aiming to disentangle the mechanisms structuring microbial communities in equatorial observatories should consider weekly or even daily sampling in relation to tidal cycles, or during critical periods such as the rainy (April–July) and windy (August–November) months in the EAMO region.

The structure of microbial co-occurrence networks further emphasized latitudinal contrasts in assembly mechanisms. The stronger dominance of protists in the EAMO network likely reflects its higher protist diversity compared to BBMO, particularly in the small-size fraction, thereby increasing their prominence and inferred associations in co-occurrence networks (Lima-Mendez et al., 2015). The EAMO network was characterized by lower connectivity but stronger positive association strength and higher assortativity by degree, implying a community with fewer but more stable interactions, potentially maintained by biotic interactions and environmental homogeneity. The higher average path length in EAMO networks suggests greater stability through slower disturbance propagation, as fewer direct connections and longer paths reduce cascading effects in more homogeneous environments (Coyte et al., 2015; Deutschmann et al., 2024). In contrast, the BBMO network exhibited higher edge density, transitivity, and domain-based assortativity, indicating more modular and cohesive microbial interactions (Deutschmann et al., 2023). These patterns may reflect both a higher number of co-occurring taxa and more predictable seasonal recurrences at BBMO, consistent with classic ecological theory (Margalef, 1978) and microbial time-series studies showing seasonal succession driven by light and temperature (Auladell et al., 2022; Caracciolo et al., 2022; Faust & Raes, 2012; Gilbert et al., 2012; Lambert et al., 2019; Yeh & Fuhrman, 2022).

The strong correlation between network metrics and seasonal variables at the BBMO, but not at the EAMO, suggests that environmental selection structures not just community composition, but also microbial interaction networks in the temperate ocean (Deutschmann et al., 2023). The EAMO network metrics, however, correlated with nitrate and silicate concentrations, reinforcing that the microbial community structure is likely being affected by non-seasonal episodic sedimentary mixing through tidal currents (Wijaya et al., 2023) in the equatorial site. For example, the positive correlation between edge density and positive strength with silicate suggest sediment resuspension could lead to a more stable and connected microbial community in the EAMO. Overall, our network and ecological analyses converge on a coherent view: tropical microbial communities, while less connected, are more stable and dominated by stochastic or biotic processes, whereas temperate communities are shaped by stronger environmental selection that also influences the structure and dynamics of microbial interactions.

We show in this study that beyond the expected differences in microbial community patterns in coastal observatories from contrasting latitudes, the ecological processes that generate these patterns are fundamentally different. Deterministic selection plays a greater role in shaping community turnover and interaction networks in the temperate site, largely due to stronger environmental gradients such as daylength and temperature. Conversely, tropical microbial communities are more temporally stable and likely more influenced by biological interaction and stochastic processes. This study further reinforces the value of comparative time-series across latitudes for disentangling the drivers of microbial community structure in the ocean. As climate change alters the amplitude and frequency of environmental fluctuations, understanding the balance between deterministic and stochastic forces in shaping microbiomes will be essential to better predict future ecosystem responses. This underscores the need for more continuous tropical time-series observatories, particularly those applying molecular tools, which remain underrepresented globally compared to temperate datasets.

## Acknowledgments

This research was supported by FAPESP (process #2014/13139-3), CNPq (process 474759/2013-0), and the Brazilian Science Without Borders Program (BJT 013/2012; PVE 400313-2014-6). PCJ was supported by Fundação de Amparo à Pesquisa do Estado de São Paulo – FAPESP (PhD grants #2017/26786-1 and #2020/02517-4) and FAI/UFSCar (ProEx n° 3213/2020-83) through the European Union – H2020 project AtlantECO (award n° 862923). HS and AMA gratefully acknowledge continuous funding through Research Productivity Grants provided by CNPq (Process: 303906/2021-9 and process: 313784/2023-0). The Blanes Bay data were obtained thanks to project DEVOTES (FP7-ENV-2012-308392), MINIME (PID2019-105775RB-I00), and MAORI (PID2022-136281NB-I00) as well as many others that have maintained the Observatory. We thank all the Blanes Bay Microbial Observatory staff, especially Vanessa Balagué, Clara Cardelús and Irene Forn for their field and laboratory support. The ICM authors acknowledge the ‘Severo Ochoa Centre of Excellence’ accreditation (CEX2024-001494-S) to the ICM-CSIC. Metabarcoding raw data processing was performed at the PIRAYU cluster (https://cimec.org.ar/c3/pirayu/index.php) via grants obtained from the Agencia Santafesina de Ciencia, Tecnología e Innovación (ASACTEI; Res N° 117/14). We thank Victor Saito and Inessa Bagatini for constructive comments on a previous version of this manuscript.

## Author contributions

PCJ, VSK, JMG and HS designed the study. HS idealized and established the EAMO. JMG idealized and established the BBMO. HS, MM, AMA, and VSK performed the EAMO fieldwork. HS, VSK, and PCJ extracted the DNA from the EAMO samples. RL, JMG, and IF sustained the long-term BBMO sampling. PCJ performed the primary data analysis and visualization. RP carried out the nutrient concentration analysis. PH assisted with the metabarcoding data processing. CRG helped with the LSP analysis, while IMD and SC helped with the network analyses. All authors assisted with interpretation of the data. PCJ, VSK, and HS wrote the manuscript. All authors contributed substantially to manuscript revisions.

## Data accessibility

The raw DNA sequences obtained in this study were deposited in the European Nucleotide Archive (http://www.ebi.ac.uk/ena) under accession number PRJEB48035 for the BBMO, and in the NCBI (https://www.ncbi.nlm.nih.gov/) under accession number PRJNA414763 for the EAMO. The ASV tables, and corresponding taxonomic classifications are deposited in Zenodo: 10.5281/zenodo.16866531 for the 16S V4V5, and 10.5281/zenodo.16866563 for the 18S V4. The contextual data is also deposited in Zenodo: 10.5281/zenodo.16861237. The processed tables and R scripts to reproduce the analyses are available in GitHub: https://github.com/pcjunger/eamo-bbmo/.

## Notes

### Competing Interest Statement

The authors have declared no competing interest.

