## Supplemental Information for "Ecological processes shaping marine microbial assemblages diverge between equatorial and temperate time-series"

**Short title:** Marine microbial assembly processes in contrasting latitudes

**Keywords:** Equatorial Atlantic, Mediterranean Sea, marine microbiome, microbial observatory, metabarcoding, network analysis

### Supplemental Information

#### Material and methods

##### *Analytical methods*

Bacterial production (BP) rates were estimated using the [3H]-leucine incorporation method (Kirchman, 1992). For this, 15 μl of [3H]-leucine (final concentration 20 nM) were added to six 1.2 ml replicates—comprising four treatments and two dead controls (to which leucine and TCA were added prior to sample addition). The samples were incubated in the dark at in situ temperature for approximately 2.5 hours. The reaction was then halted by adding 90 μl of 100% trichloroacetic acid (TCA), after which the samples were frozen at -80°C for later analysis. Bacterial proteins were extracted by washing the samples with 5% TCA (Smith & Azam, 1992), and then measured using a Beckman LS-6500 liquid scintillation counter. Disintegration rates were converted to $\mu g\ C\ l^{-1}\ h^{-1}$ using the conversion factor of 0.86 (Smith & Azam, 1992).

##### *Mock community preparation for Illumina internal control*

The mock community was prepared using near full-length amplified 16S rRNA gene clones obtained from BBMO bacterial (Alonso-Sáez et al., 2007) and archaeal (Massana et al., 2000) clone libraries. Bacterial clones were selected from different seasonal samples (Spring, Summer, and Fall; Winter samples were excluded due to unavailability of clone plates) amplified with primers 27F/1492R, ensuring broad diversity, while archaeal clones representing Euryarchaeota and Thaumarchaeota were amplified with primers 21F/958R. Selected clones were amplified using primers M13F/M13R directly from glycerol stocks (bacteria) or plasmid DNA (archaea), and the resulting PCR products were sequenced to confirm taxonomic identity and sequence quality. Clones with confirmed identity were cultured in LB medium, and plasmid DNA was extracted using a MiniPrep kit, followed by quantification using a Nanodrop spectrophotometer. The extracted plasmids were amplified with primers M13F/M13R, and the amplicons were verified by agarose gel electrophoresis to confirm expected amplicon sizes (bacterial amplicons >> archaeal amplicons). PCR products were purified using a Qiagen purification kit, quantified using a Qubit fluorometer, and tested for compatibility with sequencing primers (Parada et al., 2016) through a secondary PCR. Finally, the mock community was prepared by normalizing M13F/M13R PCR products to 20 ng/μL and combining them at a final concentration of 10 ng/μL, consistent with the sequencing facility requirements. The final set of clones included in the mock community is listed in Table 1.

**Table S1.** Final set of clones included in the mock community.

| Clone ID | Taxonomy |
|---|---|
| AUT38 | Actinobacteria - Actinomarina |
| AUT76 | Actinobacteria - Uncultured |
| AUT4 | Alphaproteobacteria - Erythrobacter |
| SPR32 | Alphaproteobacteria - Rhodobacterales |
| SPR20 | Alphaproteobacteria - SAR11 - Clade I |
| SUM5 | Alphaproteobacteria - SAR11 - Clade I |
| SUM16 | Alphaproteobacteria - SAR11 - Clade II |

| | |
|---|---|
| SUM19 | Alphaproteobacteria - SAR11 - Clade II |
| SUM1 | Alphaproteobacteria - SAR11 - Clade III |
| SUM94 | Alphaproteobacteria - SAR116 |
| AUT18 | Alphaproteobacteria - Uncultured |
| AUT45 | Alphaproteobacteria - Uncultured |
| AUT80 | Alphaproteobacteria - Uncultured |
| SUM18 | Bacteroidetes - NS5_marine_group |
| SPR33 | Bacteroidetes - NS7_marine_group |
| AUT2 | Cyanobacterium - Prochlorococcus |
| AUT89 | Cyanobacterium - Prochlorococcus MIT0801 |
| SUM10 | Cyanobacterium - Synechococcus |
| AR87 | Euryarchaeota |
| AUT3 | Alphaproteobacteria - SAR11_clade |
| AUT16 | Gammaproteobacteria - SAR86_clade |
| AUT27 | Gammaproteobacteria - SAR86_clade |
| AUT55 | Gammaproteobacteria - SAR86_clade |
| SUM93 | Gammaproteobacteria - SAR86_clade |
| ARP3 | Thaumarchaeota |
| SUM55 | Verrucomicrobia |
| AUT17 | Verrucomicrobia - Pelagicoccus |

##### *DNA sequencing*

The DNA PCR amplification and sequencing were conducted at the Integrated Microbiome Resource (IMR, Dalhousie University, Halifax, Canada; http://imr.bio/index.html) for the 16S rRNA, and at the Functional Genomics Centre (ESALQ-USP, University of São Paulo, Piracicaba-SP, Brazil; https://sites.usp.br/cgf/) for the 18S rRNA. The 16S rRNA PCR products were normalized and purified with the Charm Biotech Just-a-Plate Purification and Normalization kit, following the sequencing facility protocol (Comeau & Kwawukume, 2023).

##### *Determining seasonal ASVs*

Function *randlps()* computes the Lomb-Scargle periodogram and the p-values for the largest peak in the periodogram by randomising the time-series sequence (Ruf, 1999). The choice of the type of normalization — "standard" or "press" — determines the values of the periodogram peaks, which are confined to the interval 0-1 if normalization = "standard", or normalized using the factor 1/(2 * var(y)) if normalization = "press". Studies using the LPS approach with normalization = "standard" set the threshold at PNmax > 0.1 (Jing et al., 2024; Zhao et al., 2023), and studies with normalization = "press" set it at PNmax >10 (Auladell et al., 2022; Ferrera et al., 2024; Lambert et al., 2019).

We decided to use the default parameters in the *randlps()* function with normalization = "standard". We manually plotted the rarefied abundance of all ASVs with PNmax > 0.1 and $p < 0.01$ across months and years, to inspect the seasonality trend (see example below with *Bathycoccus prasinos* 0.22-3 μm with PNmax = 0.35, $p < 0.001$) and decided to set the threshold to 0.2 to keep only robust signals of seasonality. Finally, since the function looks for all possible rhythmic patterns in a signal, regardless of their period, we also checked that all selected ASVs showed a period of ~1 year.

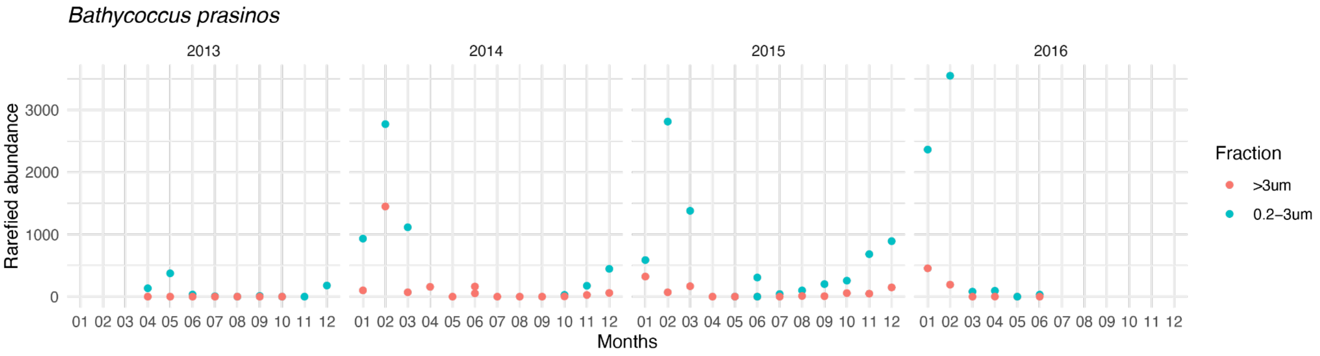

*Network construction and filtering*

**Table S2.** Summary of the sample and ASV filtering for network construction

| Location | | **BBMO** | | | | **EAMO** | | | |
|---|---|---|---|---|---|---|---|---|---|
| Kingdom | | **16S** | | **18S** | | **16S** | | **18S** | |
| Size fraction | | **0.22-3** | **>3** | **0.22-3** | **>3** | **0.22-3** | **>3** | **0.22-3** | **>3** |
| **Sample Filtering** *Samples need to contain both kingdoms (16S and 18S) and both size fractions (0.22-3 µm and >3 µm)* | | | | | | | | | |
| #Samples | | 40 | 30 | 33 | 39 | 30 | 22 | 29 | 29 |
| Both size-fractions within kingdom | | 29 | | 31 | | 22 | | 27 | |
| Both size fractions and both kingdoms | | 23 | | | | 22 | | | |
| **#ASV filtering done for each table (before Sample Filtering)** *ASVs need to have an abundance sum above 100 counts and be present in more than 15% of samples* | | | | | | | | | |
| Abundance | | 1137 | 578 | 2547 | 2395 | 831 | 308 | 2358 | 2024 |
| Prevalence | | 981 | 617 | 662 | 619 | 924 | 507 | 1657 | 1220 |
| Both filters | | 818 | 442 | 628 | 598 | 655 | 257 | 1447 | 1083 |
| **#ASV filtering done for samples containing both size fractions and both kingdoms** | | | | | | | | | |

| | | | | | | | | | |
|---|---|---|---|---|---|---|---|---|---|
| Both filters | | 797 | 414 | 577 | 690 | 578 | 257 | 1296 | 1114 |
| **Size Fraction Filtering**<br>*Size fraction filtering: for each ASV, if the ratio of counts (`big size / small size`) is less than 0.5, the ASV is removed from the big size fraction. Similarly, if the ratio is greater than 2, the ASV is removed from the small size fraction.* | | | | | | | | | |
| #ASV removed | | 19 | 239 | 43 | 146 | 14 | 117 | 208 | 339 |
| #ASV remaining | | 778 | 175 | 534 | 544 | 564 | 140 | 1088 | 775 |

**Table S3.** Summary of the network filtering with EnDED

| Network | | **BBMO** | | | | **EAMO** | | | |
|---|---|---|---|---|---|---|---|---|---|
| Kingdom | | **16S** | | **18S** | | **16S** | | **18S** | |
| Size fraction | | **0.22-3** | **3-200** | **0.22-3** | **3-200** | **0.22-3** | **3-200** | **0.22-3** | **3-200** |
| #ASV | | 778 | 175 | 534 | 544 | 564 | 140 | 1088 | 775 |
| #ASV (all) | | 2031 | | | | 2567 | | | |
| **Network constructed with FlashWeave**<br>*Only nodes with at least one edge are considered, i.e., isolated nodes are removed.* | | | | | | | | | |
| #nodes | | 756 | 159 | 468 | 531 | 413 | 87 | 920 | 679 |
| #nodes (all) | | 1914 | | | | 2099 | | | |
| #edges | | 2660 (2395 pos, 265 neg) | | | | 2435 (2073 pos, 362 neg) | | | |
| **EnDED** | | | | | | | | | |
| #edges removed | | 6 (2 pos, 4 neg)<br>daylength: 0<br>Temperature: 0<br>Salinity: 0<br>NH4: 0<br>NO2: 3<br>NO3: 1<br>PO4: 1<br>Si: 1 | | | | 14 (5 pos, 9 neg)<br>daylength: 0<br>Temperature: 0<br>Salinity: 0<br>NH4: 4<br>NO2: 2<br>NO3: 2<br>PO4: 1<br>Si: 6 | | | |
| #edges | | 2654 (2393 pos, 261 neg) | | | | 2421 (2068 pos, 353 neg) | | | |
| #nodes | | 1913 | | | | 2091 | | | |
| **EnDED: filtering based on percentage co occurrence > 50%** | | | | | | | | | |

| | | | |
|---|---|---|---|
| #edges removed | | 40 (0 pos, 40 neg) | 32 (0 pos,32 neg) |
| #edges | | 2614 (2393 pos, 221 neg) | 2389 (2068 pos, 321 neg) |
| #nodes | | 1906 | 2074 |

#### Supplementary Figures

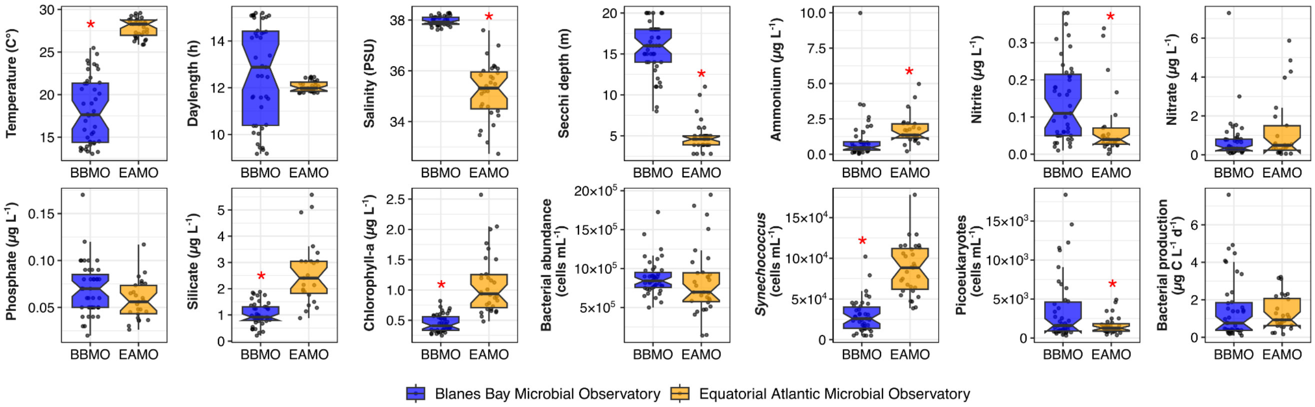

**Figure S1.** Comparison of environmental and biological variables between the two observatories (BBMO – Temperate site; EAMO – Tropical site). Red asterisks indicate significant statistical differences (t-test, $p<0.01$).

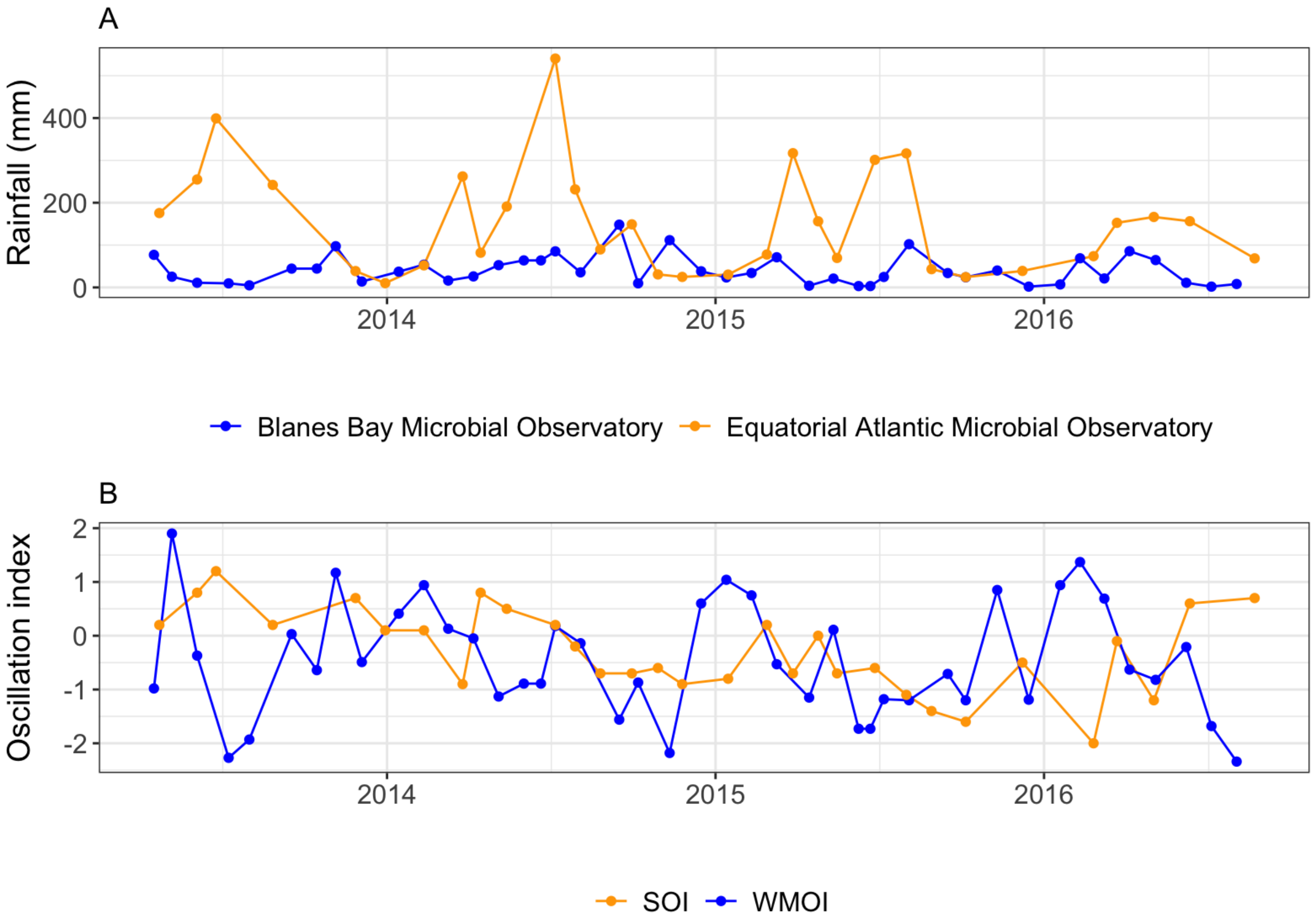

**Figure S2.** Time-series (April 2013 to August 2016) of the additional climatic and meteorological variables obtained from public databases, as described in the methods sections. **(A)** Monthly accumulated rainfall prior to the sampling date; **(B)** Monthly values of the Southern Oscillation (SOI) and the Western Mediterranean Oscillation (WMOI) indexes.

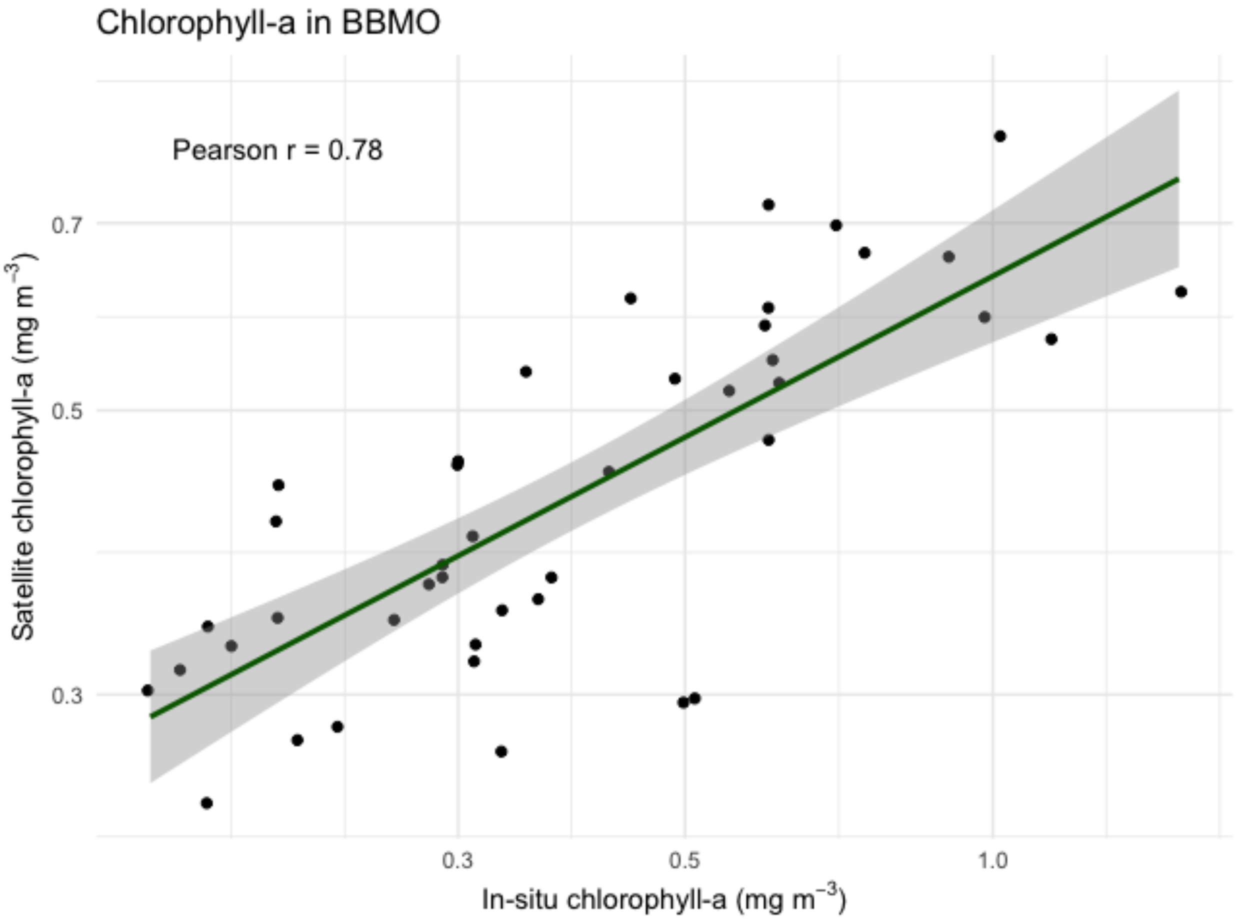

**Figure S3.** Correlation between satellite-derived and *in situ* chlorophyll *a* estimates at the Blanes Bay Microbial Observatory (BBMO). The correlation was statistically significant (Pearson's r= 0.78, $p<0.01$).

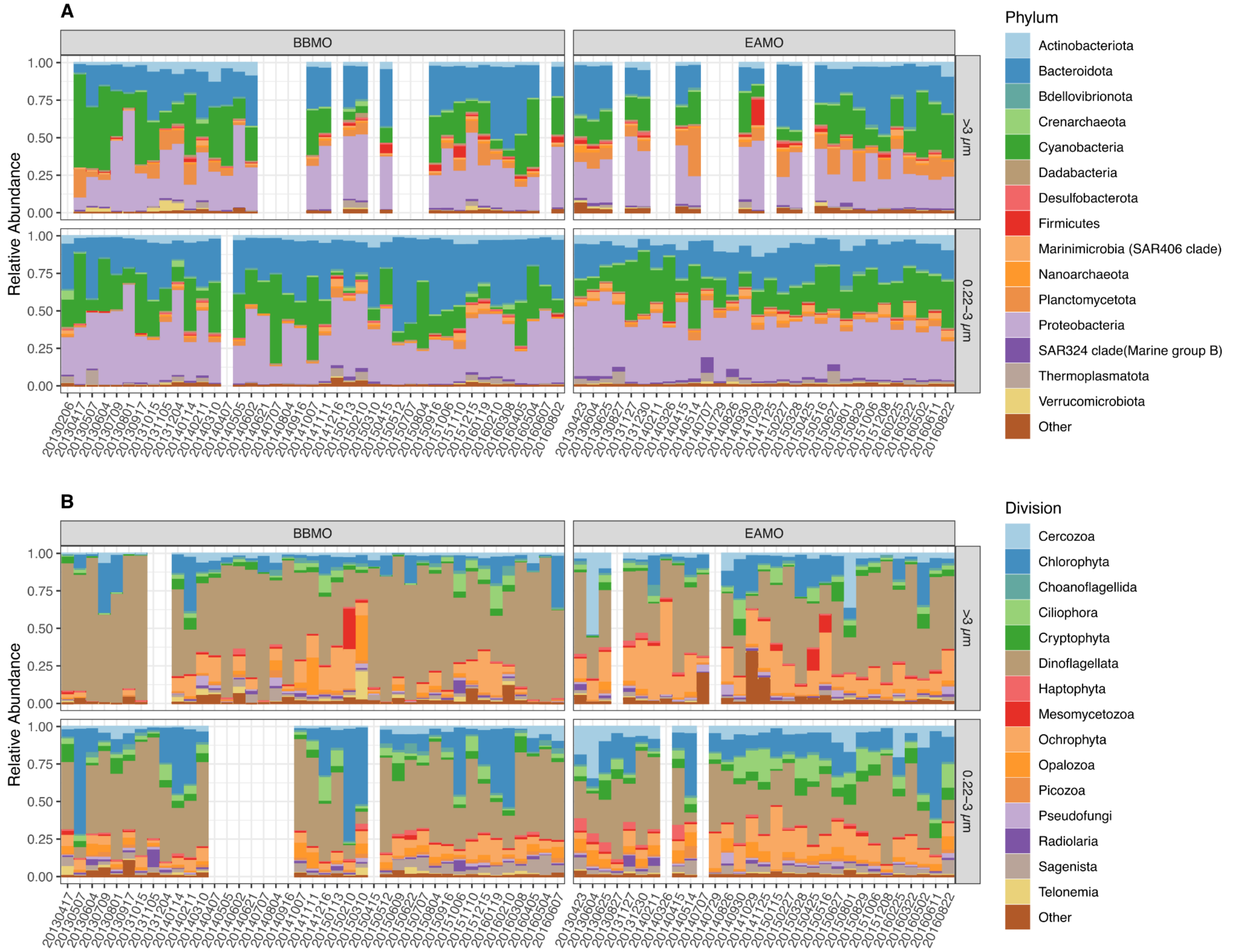

**Figure S4.** Taxonomic composition across samples of the **A)** prokaryotic and **B)** protist community in both size-fractions (0.22-3 µm and >3 µm) at the Blanes Bay Microbial Observatory (BBMO – Temperate site) and the Equatorial Atlantic Microbial Observatory (EAMO – Tropical site). The empty columns are samples with sub-optimal sequencing.

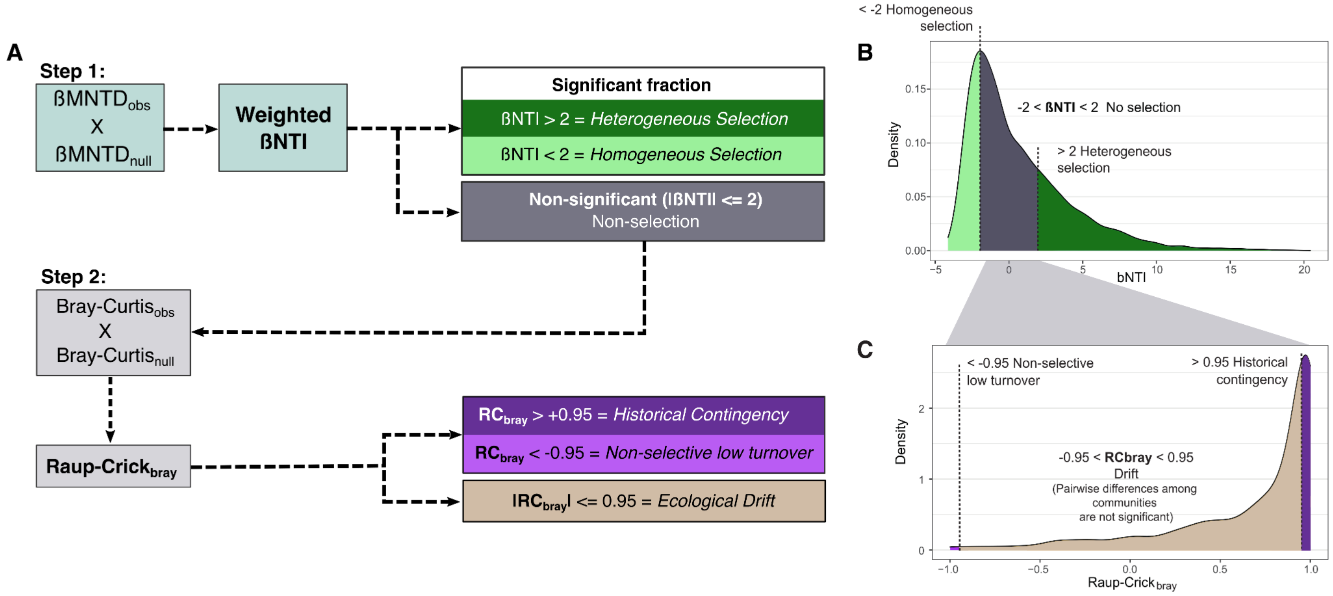

**Figure S5. A)** Summary of the analytical framework adapted from Stegen et al. 2013 to estimate ecological processes from probabilistic models using time-series data. The visualization of the model distributions **(B, C)** is adapted from (Gazulla et al., 2022). **B)** Distribution of βNTI distribution across all pairwise comparisons in the dataset. Absolute βNTI values greater than 2 indicate significant deviations from random phylogenetic turnover, suggesting the influence of either homogeneous or heterogeneous selection. The grey area highlights the range of nonsignificant βNTI values. To further differentiate whether drift or historical contingency are driving community turnover among these comparisons, we calculated the Bray–Curtis-based Raup–Crick metric ($RC_{bray}$). **C)** Distribution of $RC_{bray}$ values for the subset of pairwise comparisons not structured by selection. $RC_{bray}$ values between –0.95 and +0.95 suggest community assembly dominated by drift. Values > +0.95 or < –0.95 indicate that turnover is primarily shaped by historical contingency or rare non-selective processes, respectively.

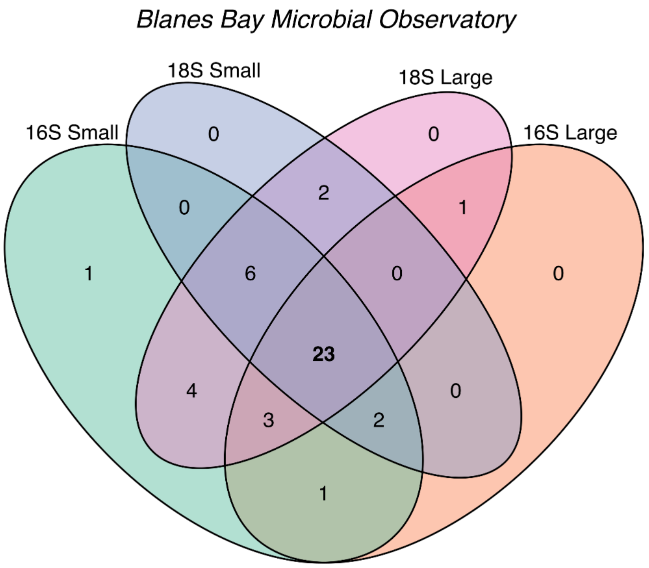

**Figure S6.** Venn's diagrams show the number of samples that had both 16S (prokaryotes) and 18S (protists) data, and "small" (0.22–3 µm) and "large" (>3 µm) size fractions. These samples were therefore retained for the network construction of BBMO (n=23) and EAMO (n=22).

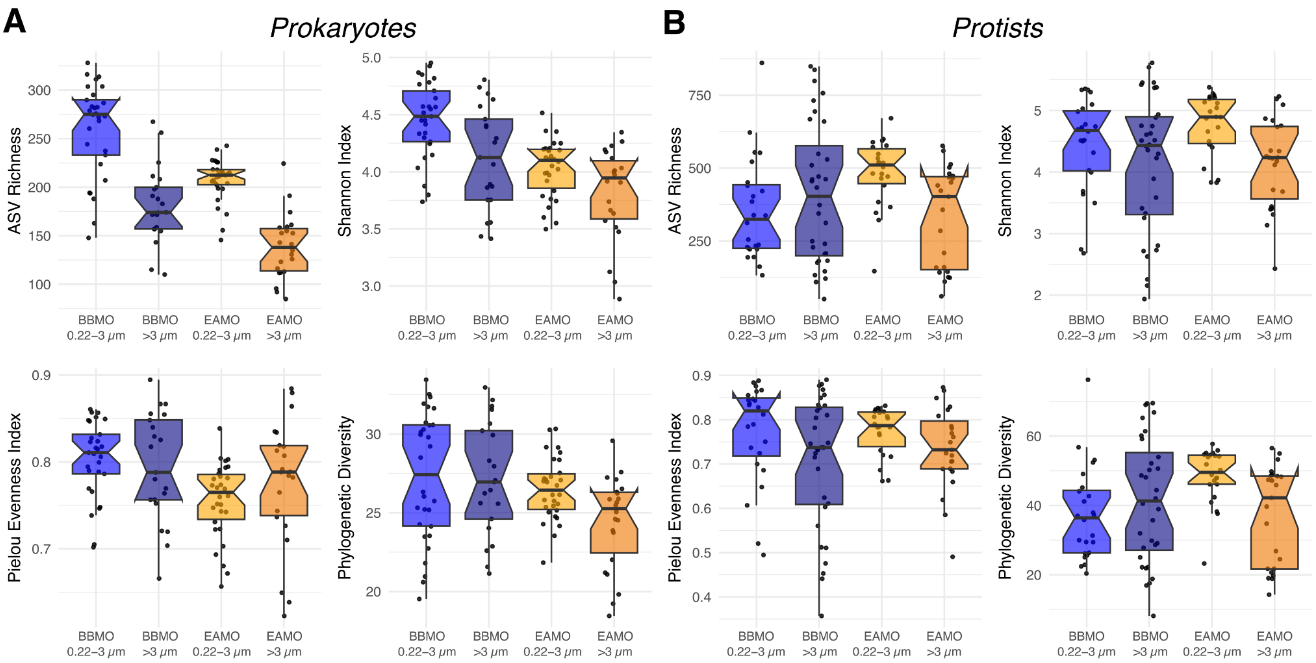

**Figure S7.** Diversity metrics showing the ASV richness, Pielou's evenness index, Shannon diversity index, and phylogenetic diversity, for each site and fraction.

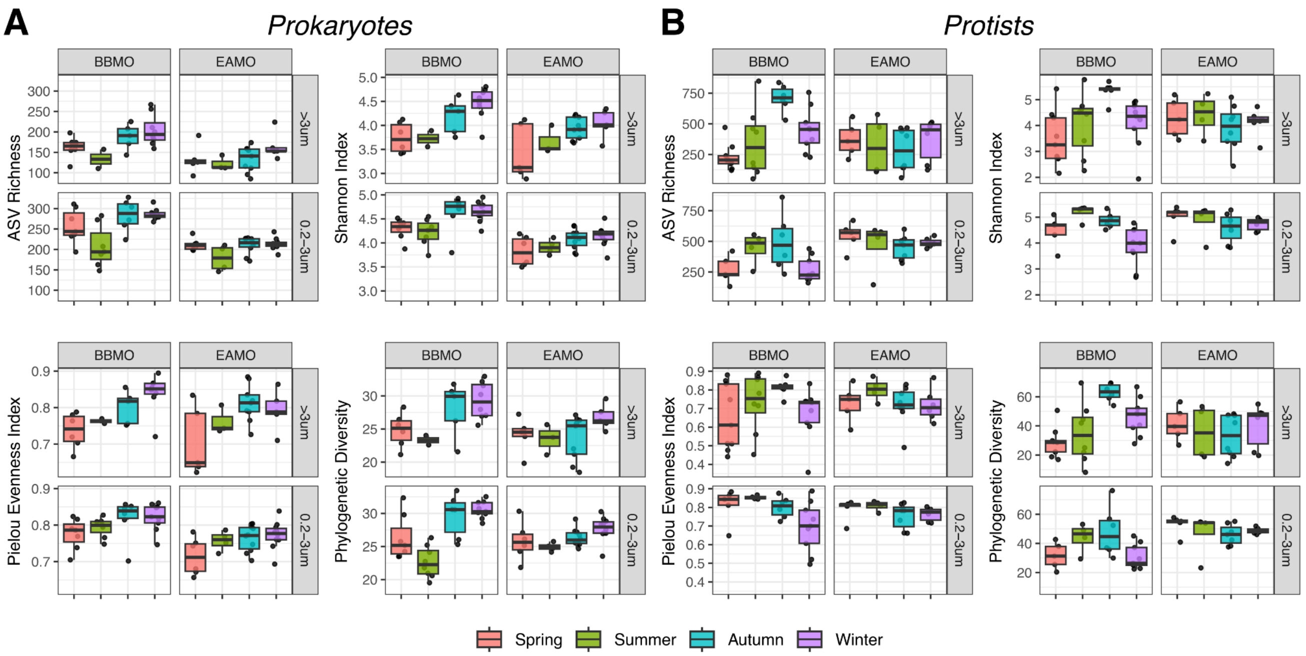

**Figure S8.** Seasonal differences in diversity metrics of **(A)** prokaryotes and **(B)** protists in the Blanes Bay Microbial Observatory (BBMO) and the Equatorial Atlantic Microbial Observatory (EAMO). The seasons were astronomically defined based on the dates in the Northern and the Southern hemispheres, respectively.

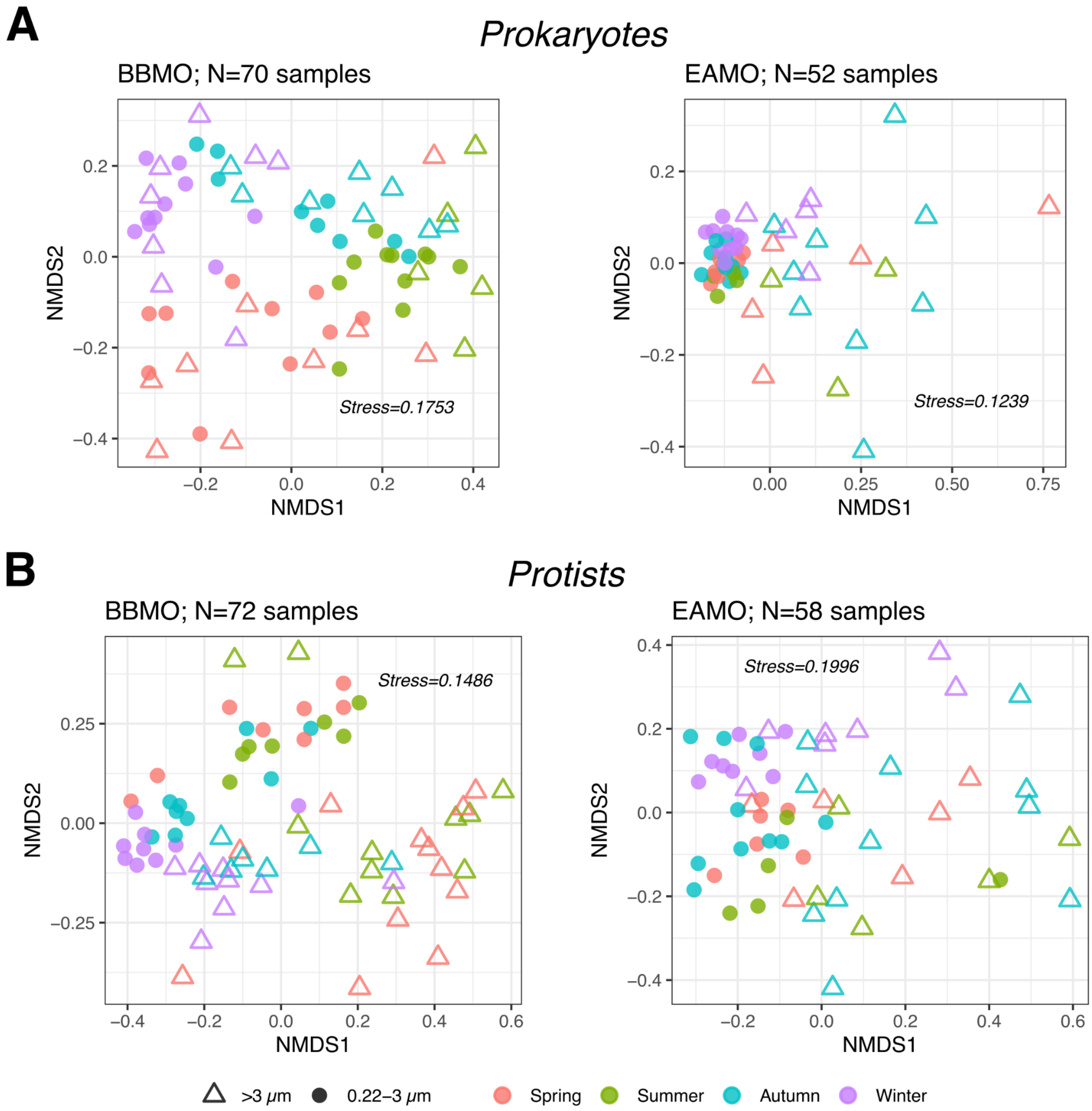

**Figure S9.** Nonmetric multidimensional scaling (NMDS) based on the Bray-Curtis dissimilarities among prokaryotic and eukaryotic samples – labeled by seasons (colors) and size-fraction (shapes).

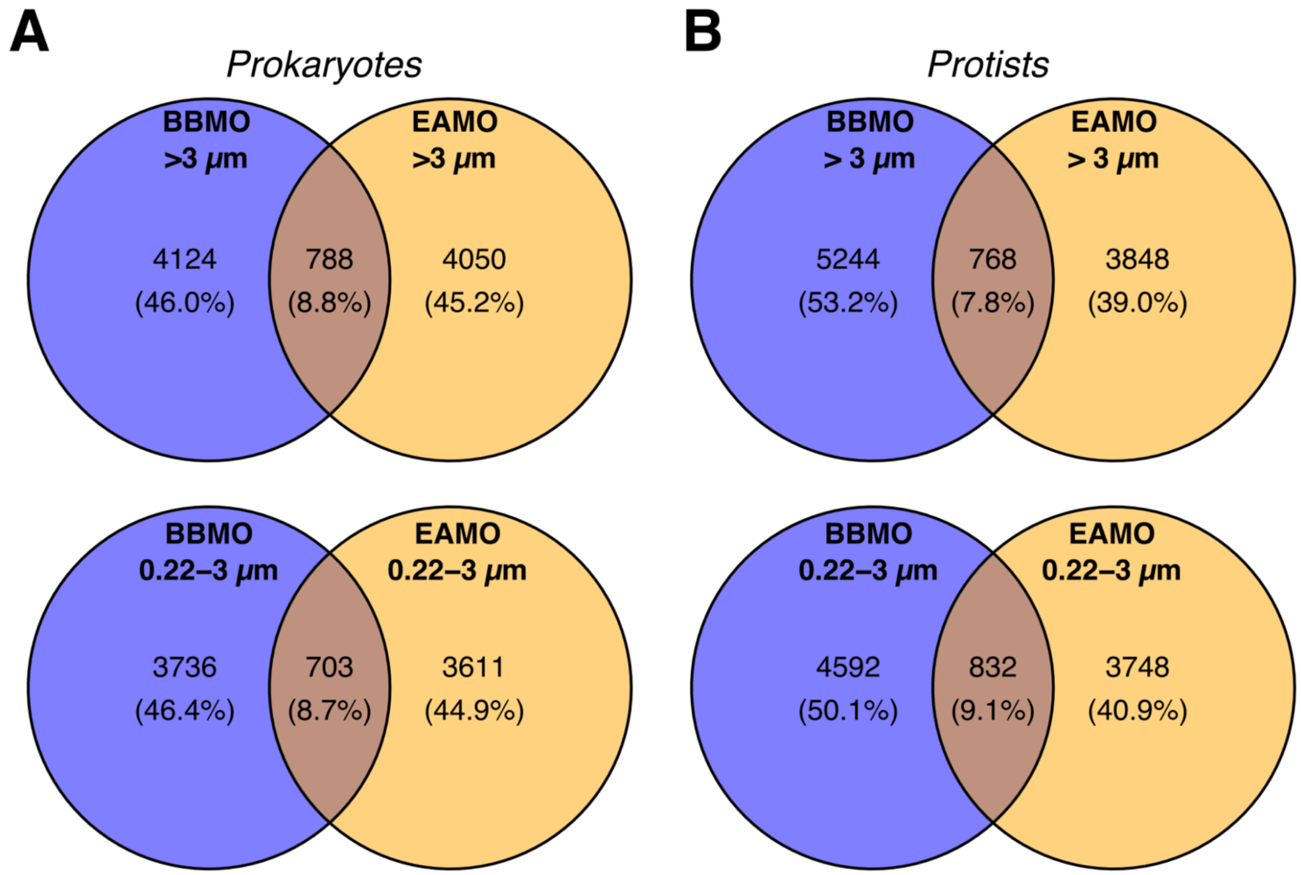

**Figure S10.**Venn's plots with the number of common and unique **(A)** prokaryotic and **(B)** protist ASVs in each size-fraction between sites.

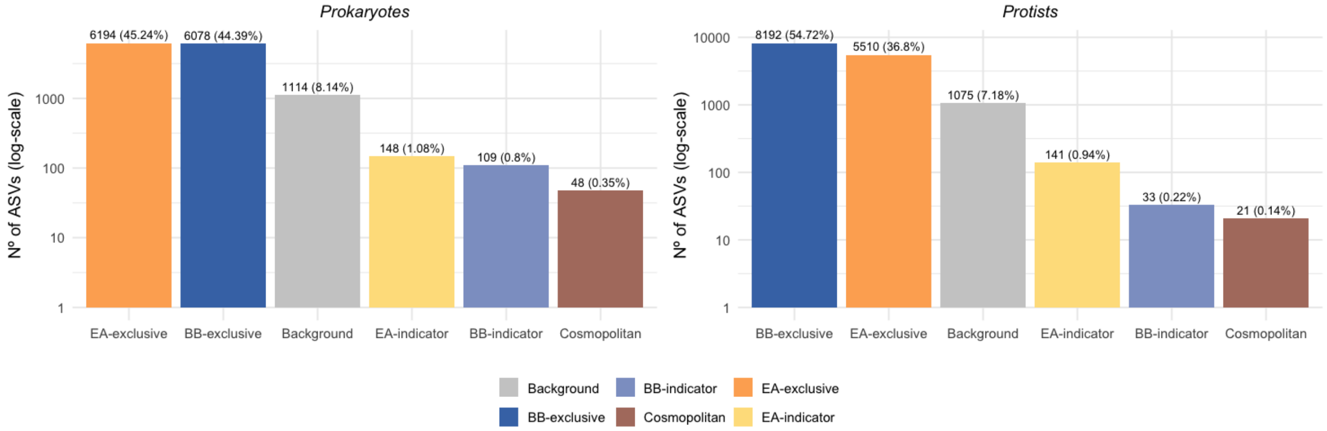

**Figure S11.** Number of ASVs within each category, as described in the methods. The percentage of total ASVs is indicated above each bar. Bars are sorted in decreasing order by number of ASVs. BB – Blanes Bay Microbial Observatory; EA – Equatorial Atlantic Microbial Observatory.

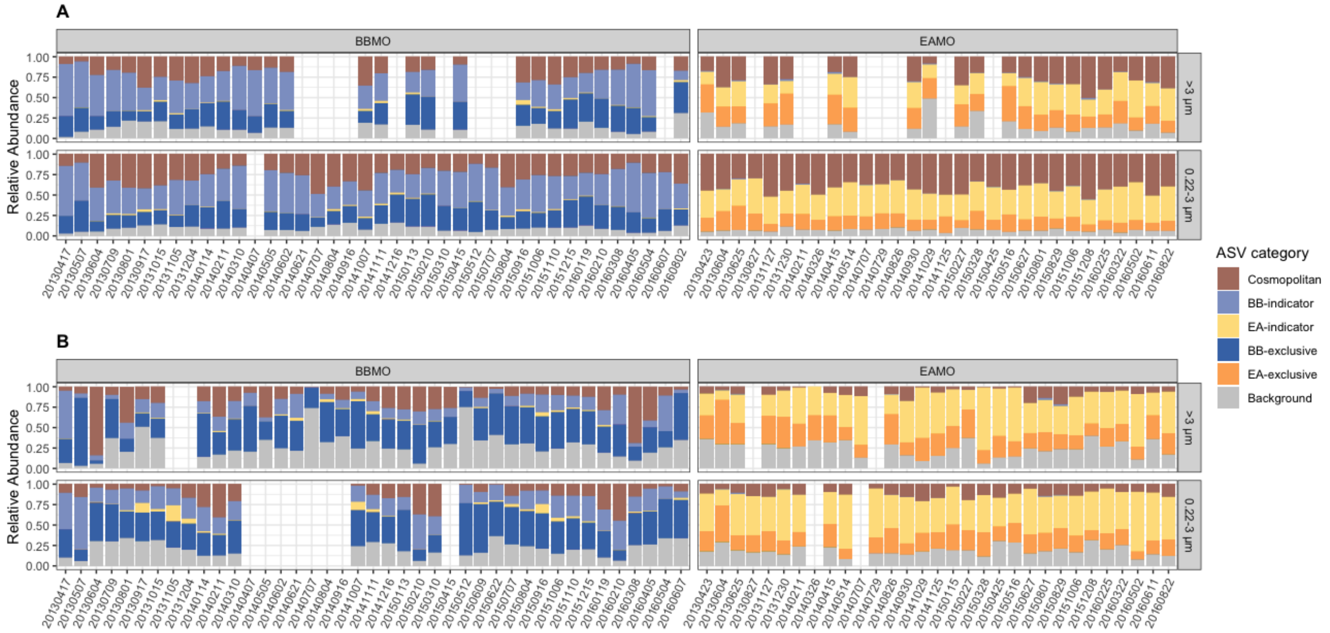

**Figure S12.** Relative abundance of the ASV categories of the **(A)** prokaryotic and **(B)** protist communities across samples. BBMO – Blanes Bay Microbial Observatory; EAMO – Equatorial Atlantic Microbial Observatory. The empty columns are samples with sub-optimal sequencing.

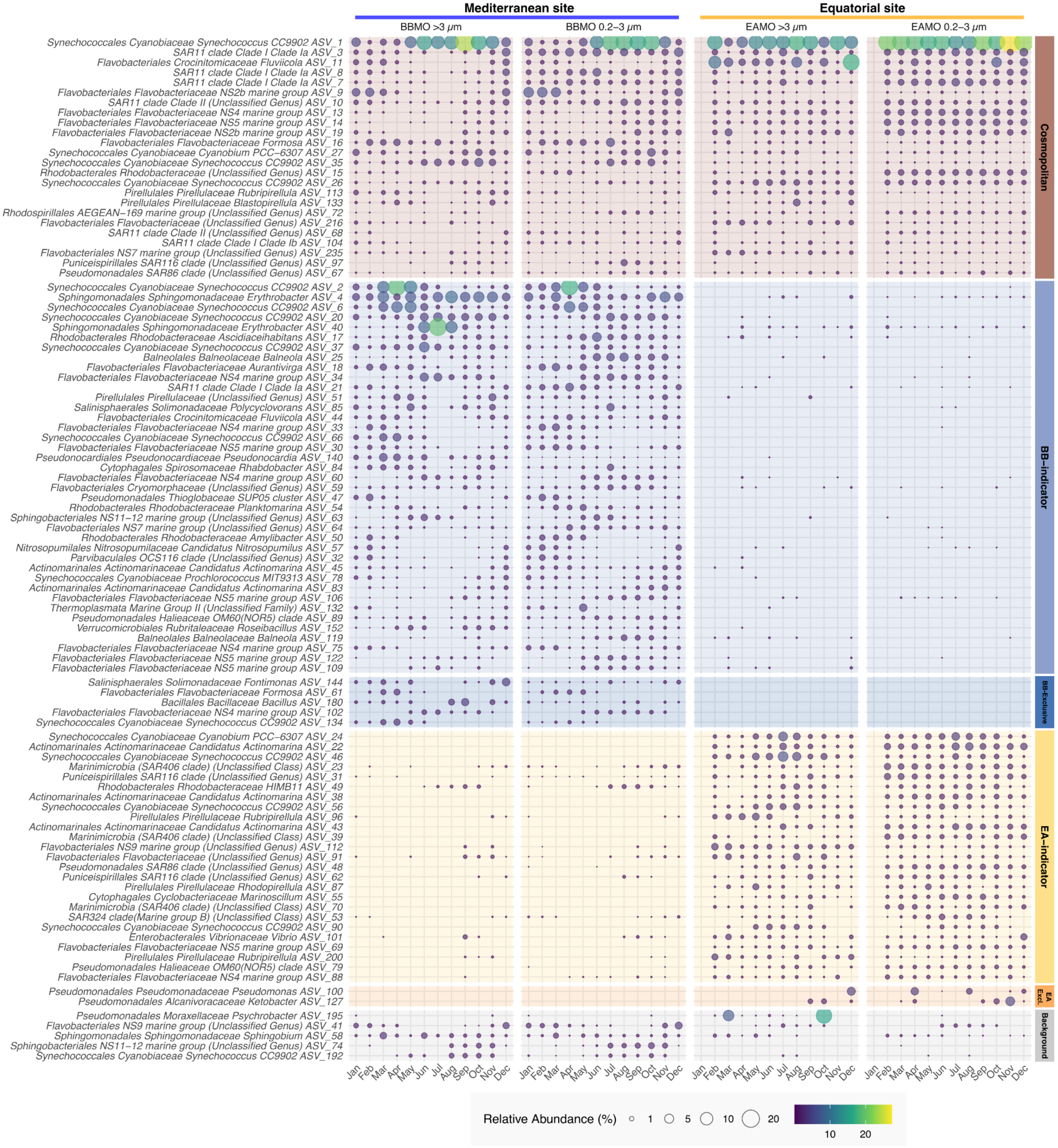

**Figure S13.** The monthly average relative abundance of the 100 most abundant prokaryotic ASVs classified as cosmopolitan, BB-indicators, EA-indicators, BBMO-exclusive, EAMO-exclusive, or background. BB = BBMO, EA = EAMO. The BBMO-exclusive and EAMO-exclusive categories refer to the ASVs which were unique to the station, but not statistically determined as an indicator.

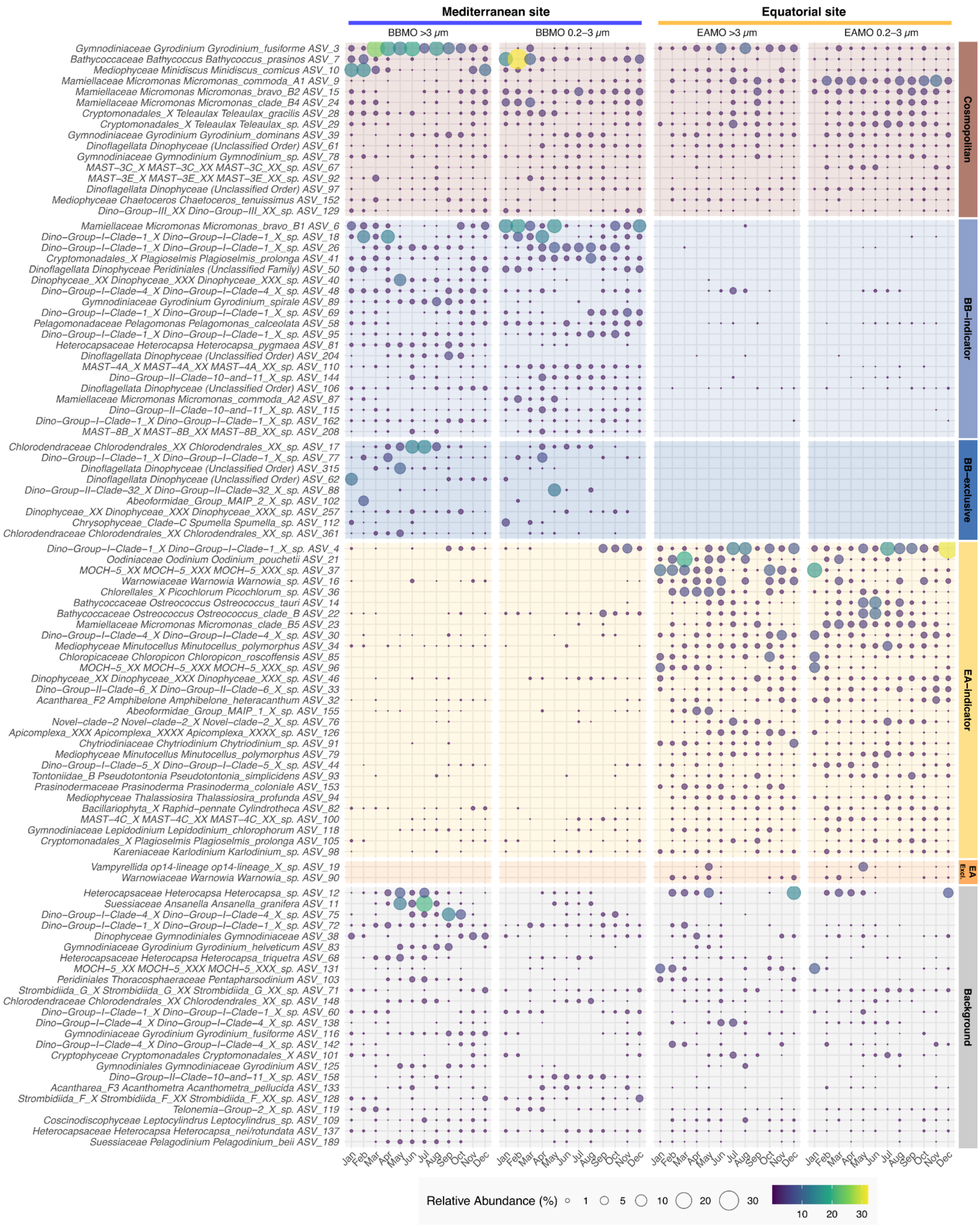

**Figure S14.** The monthly average relative abundance of the 100 most abundant protist ASVs classified as cosmopolitan, BB-indicators, EA-indicators, BBMO-exclusive, EAMO-exclusive, or background. BB = BBMO, EA

= EAMO. The BBMO-exclusive and EAMO-exclusive categories refer to the ASVs, which were unique to the station, but not statistically determined as an indicator.

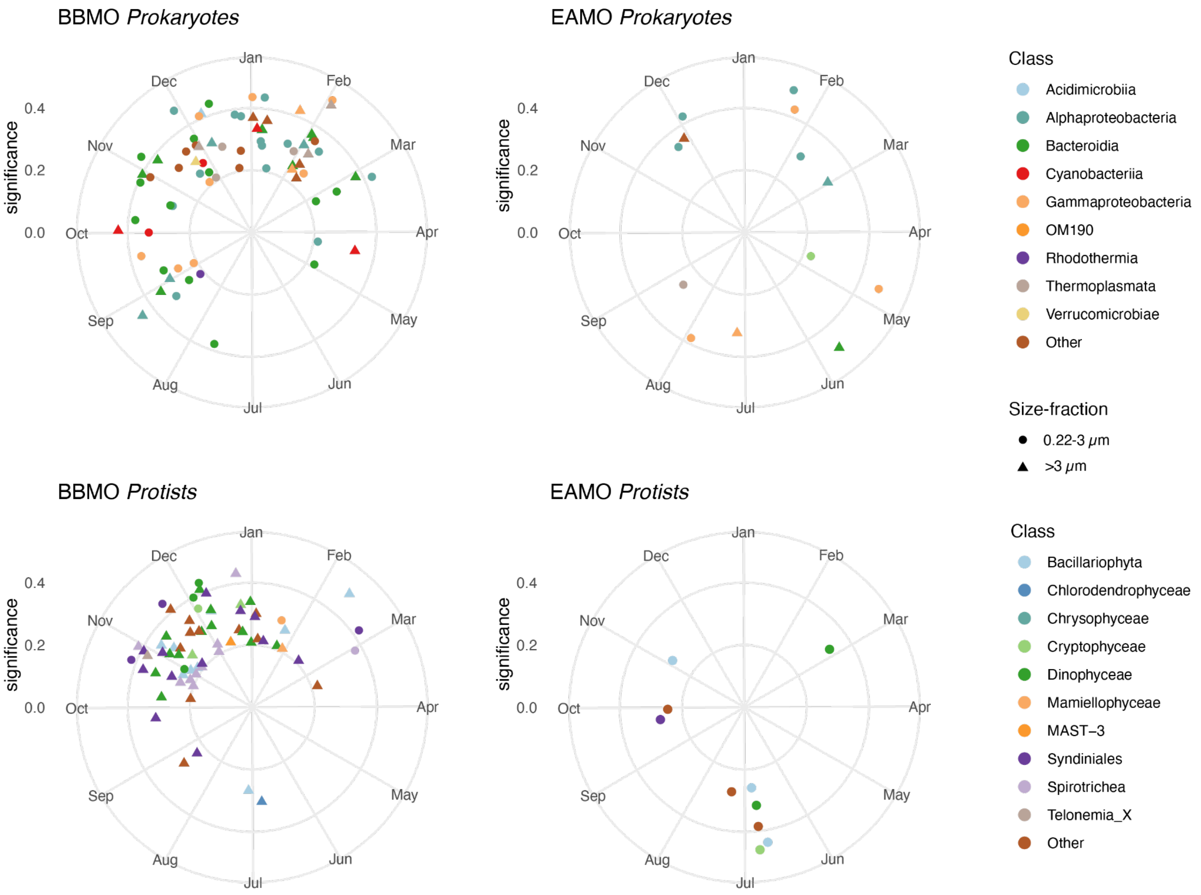

**Figure S15.** Polar plots representing the seasonal ASVs of the prokaryotic and protist communities. Different symbols indicate size fractions. ASVs are color-coded by taxonomic groups. Higher strength recurrence values represent stronger seasonal signals.

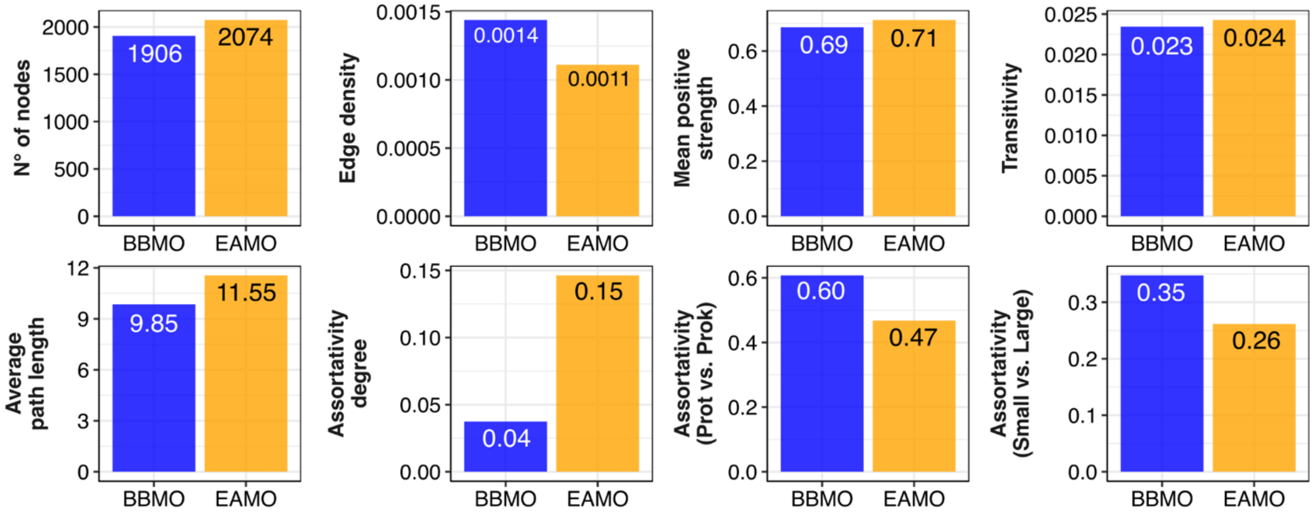

**Figure S16.** Bar plots showing the topological metrics of the Blanes Bay Microbial Observatory (BBMO) and the Equatorial Atlantic Microbial Observatory (EAMO) obtained from the static networks.
